# *Plasmodium falciparum* Niemann-Pick Type C1-Related protein relies on its physicochemical properties for membrane contact site localization required for cholesterol homeostasis

**DOI:** 10.64898/2026.03.25.714277

**Authors:** Ananya Ray, Eva. S. Istvan, Daniel E. Goldberg, Matthias Garten

## Abstract

The Niemann-Pick Type C1-Related protein of the malaria parasite *Plasmodium falciparum*, PfNCR1, is a promising anti-malarial drug target facilitating cholesterol homeostasis at the interface of the malaria parasite with its host-red blood cell. PfNCR1 is localized to otherwise functionally uncharacterized regions covering ∼half of the host-parasite interface. These regions are defined by exceptionally narrow membrane contact sites, leaving only ∼3-4 nm vertical aqueous space in between the membranes. Determining the origin and functional consequence of localization to the closely apposed membrane is central for our understanding of PfNCR1 as drug target but also offers a window into the mechanism of the group of homologous proteins, associated with congenital conditions and cancer. Here we define the mechanism of PfNCR1’s membrane contact site localization and its implication for cholesterol transport. We identified a 141 amino acid long (amphipathic) helix - linker - (amphipathic) helix domain (“HLH domain”) unique to *Plasmodium spp.*, that is necessary for efficient localization of PfNCR1 to the narrow membrane contact sites. Mechanistically, we show that this localization relies on the HLH domain’s physicochemical properties. GPI-anchoring the isolated HLH domain or a version of the HLH domain in which the helices are replaced by the amphipathic helix of human ATG3 are sufficient to target a fluorescent protein to regions of endogenous PfNCR1. Functionally, we demonstrate that the degree of PfNCR1 localization to narrow contact sites qualitatively correlates with its ability to maintain cholesterol homeostasis, linking PfNCR1’s membrane contact to its recognized transport function. Collectively, the results establish the HLH domain as key element for PfNCR1’s localization and effectiveness in cholesterol transport while also opening avenues to probe the narrow membrane contact site regions with engineered proteins.

## Introduction

Cholesterol homeostasis is required for animals and many microbes to set key membrane properties such as membrane rigidity, thickness, permeability and curvature; provide a local environment to facilitate protein-protein interaction in membranes; or serve as signal required for cell differentiation (Kong et al. 2019; Luo et al. 2020; Levental and Lyman 2023). In the malaria parasite the *Plasmodium falciparum* Niemann-Pick Type C1-Related protein, PfNCR1 (UniProt id Q8I266, PF3D7_0107500), was identified as an essential, druggable protein responsible for maintenance of low cholesterol in the parasite plasma membrane (PPM) (Istvan et al. 2019; Ahiya et al. 2022). Three compounds were found to potently inhibit PfNCR1 function. Therefore, the protein is of prime interest to target cholesterol homeostasis and has the potential to complement front line anti-malarials. This is significant for the 282 million people affected by malaria and 610,000 deaths each year, most of whom are young children (World Health Organization 2025). Mechanistic and structural insight into PfNCR1’s function using recombinant protein corroborated PfNCR1’s direct role in cholesterol homeostasis, demonstrating cholesterol transport function powered by a pH gradient (Zhang et al. 2024). Sequence similarity and the cryo-EM structure (Zhang et al. 2024) suggest that PfNCR1 is a homolog of the human Niemann-Pick disease type C intracellular cholesterol transporter 1 (hNPC1, UniProt id O15118) and human Protein patched homolog 1 (hPTCH1, UniProt id H3BLX7), notable for their link to neurodegenerative disorders due to defective cholesterol trafficking and cancer due to defective signaling in the Hedgehog pathway, respectively. While PfNCR1 was named after hNPC1, the lack of cholesterol receiving/binding domain A of hNCP1 indicates that PfNCR1’s role in maintaining plasma membrane cholesterol might be more similar to the non-Hedgehog function attributed to hPTCH1, and best characterized for patched homolog 3 (PTC-3, UniProt id H2L0G5) of *C. elegans* (Cadena del Castillo et al. 2021). Hence PfNCR1 appears as a “patched-like” protein, however it is unknown how PfNCR1 transports cholesterol in its cellular context. Insight into PfNCR1’s in situ mechanism of cholesterol trafficking is equally significant for its position as a promising anti-malarial drug target and role as a homolog of hNPC1/hPTCH1.

A distinguishing feature of PfNCR1 is its localization at remarkably narrow membrane contact sites between the parasite plasma membrane (PPM) and parasitophorous vacuole membrane (PVM) that alternate with wider spaces populated by protein export machinery (Garten et al. 2020). The contact sites measure ∼9 nm from membrane center to membrane center with ∼3-4 nm aqueous space, in contrast to 20-40 nm distances for the wider regions. Both narrow and wide regions span µm^2^-sized patches around the cell, both regions qualify as membrane contact sites, as the GPI-anchored PPM resident P113 spans the PV and connects to the PVM-resident translocon found in the wider regions (Bullen et al. 2022). This geometry is structurally similar to an immunological synapse where height differences in the contact site-participating proteins drive receptor clustering (Schmid et al. 2016; Al-Aghbar et al. 2022). Intriguingly, the narrow membrane contact sites persist in absence of PfNCR1 (Garten et al. 2020) pointing to a common feature of membrane contact sites arising from multiple proteins participating in the formation of the membrane contact site (Scorrano et al. 2019; Voeltz et al. 2024). However, for *Plasmodium* it is neither known which other proteins are localized to the narrow membrane contact site region nor what functions they may fulfill. Understanding the targeting mechanism of PfNCR1 will allow characterization of the narrow membrane contact site regions of *Plasmodium falciparum* to identify participating proteins and define new, essential, potentially druggable functions that are dependent on close membrane contact.

PfNCR1’s localization in combination with available functional data suggests that PfNCR1 may work as a short hydrophobic conduit to pump cholesterol from the PPM directly into the PVM (for example generally reviewed in (Voeltz et al. 2024)). Alternatively, more intricate mechanisms of cholesterol transport are conceivable such as a combination of Sonic Hedgehog-like sensing of cholesterol in the PVM triggering cholesterol flip from the outer to the inner leaflet of the PPM (Kinnebrew et al. 2021). Curiously, PfNCR1’s localization evokes parallels to the pool of hPTCH1 at the closely opposing membranes in the ciliary pocket, where hPTCH1 is hypothesized to aid depletion of cholesterol in its plasma (cilia) membrane (Kong et al. 2019). Identifying and altering the targeting mechanism for PfNCR1 is a crucial step defining the functional role of contact site localization in general and cholesterol homeostasis in *Plasmodium*-infected red blood cells in particular.

The aim of this work is to gain insight into PfNCR1 targeting at the host-parasite interface of *Plasmodium falciparum*. Here we propose that a predicted (amphipathic) helix - linker - (amphipathic) helix (HLH) domain facing the PV-lumen side of PfNCR1, unique to *Plasmodium spp.,* is responsible for PfNCR1’s exclusive localization to the narrow membrane contact sites in between the PVM and PPM. Progressive truncations show that deletion of the whole domain randomizes PfNCR1 localization at the host-parasite interface, correlating with dysregulation of cholesterol homeostasis. We then go on to show that the HLH domain in its entirety is sufficient to target a fluorophore into the narrow contact site regions if the protein is anchored to the PPM. Lastly, we demonstrate that targeting to the narrow membrane contact site regions of *Plasmodium falciparum* does not require a *Plasmodium*-specific helix sequence but can be reproduced replacing the amphipathic helices of the HLH domain with the amphipathic helix of human ATG3.

## Results

### A *Plasmodium*-specific domain in PfNCR1 is essential for membrane contact site targeting

To define possible features of PfNCR1 that could localize the protein to the narrow membrane contact site regions, we investigated PfNCR1’s amino acid sequence and structural features. When examining the AlphaFold2 structure prediction of PfNCR1 we noticed the pseudosymmetry of the two PV-facing domains PV1 and PV2 (as defined in (Zhang et al. 2024)) but also a unique *Plasmodium* domain (defined as amino acids (aa) 142-282) only partially resolved in (Zhang et al. 2024) presumably due to its flexible nature (Fig. 1A, SI Fig. S1). AlphaFold2 suggests that the region consists of two α-helices bridged by a disordered linker. Sequence alignment revealed that the aa142-282 region is consistently present across *Plasmodium* species but absent from homologous proteins in other apicomplexan parasites and model eukaryotes, indicating a species-specific role that may relate to its localization (SI Fig. S1).

**Fig. 1.**
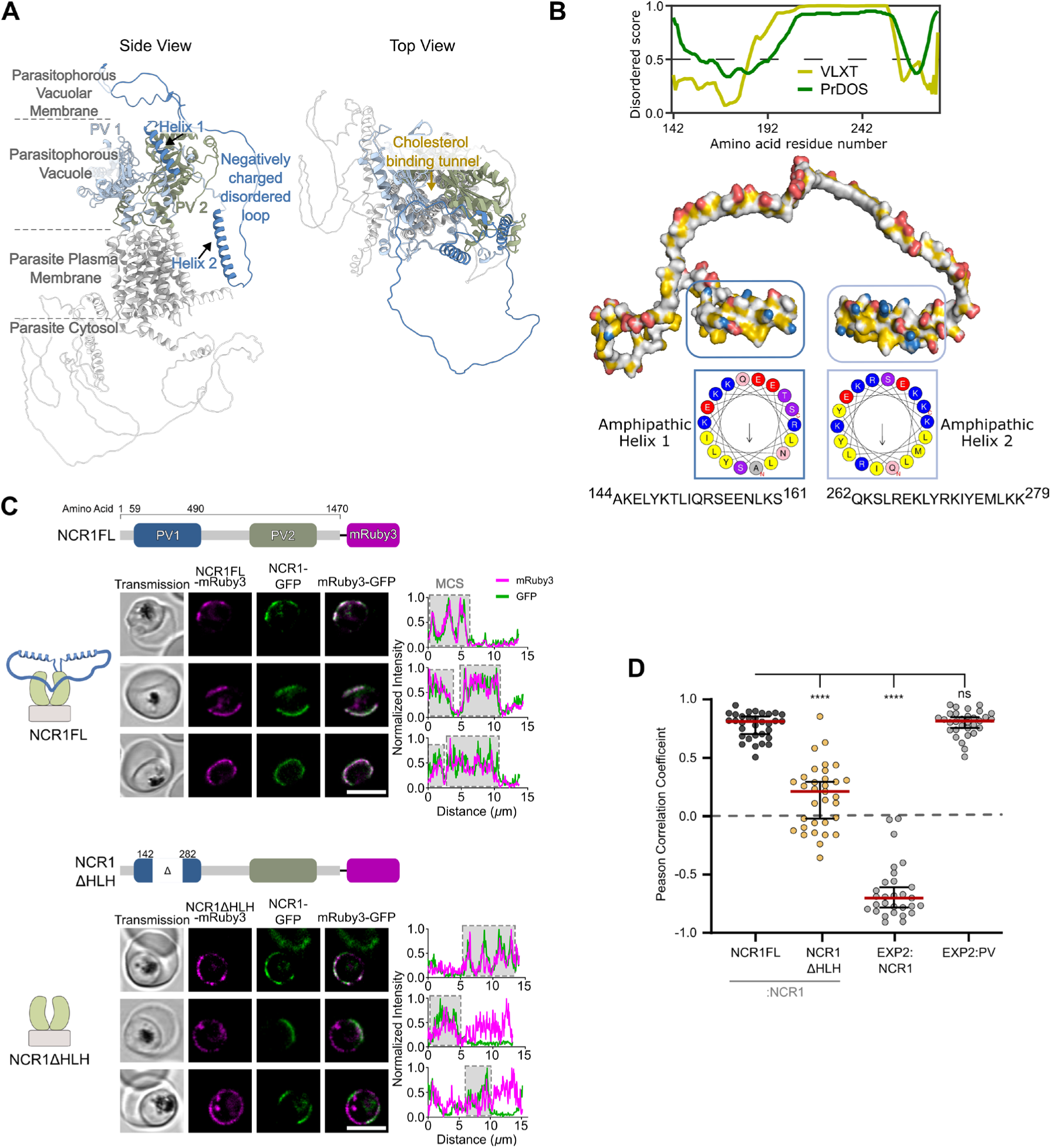
NCR1 Domain unique to *Plasmodium* is necessary for targeting to the contact sites. A. AlphaFold2 prediction of PfNCR1. Light blue and green are the two domains facing the parasitophorous vacuole, darker blue labels the HLH domain. Approximate location of the parasite plasma membrane (PPM), parasitophorous vacuole membrane (PVM), and the parasitophorous vacuole (PV) space are indicated. Top view showing the spatial positioning of the PV-facing domains around the cholesterol transport tunnel. B. Properties of the HLH domain. Disorder prediction of the HLH-domain using 2 different algorithms (VL-XT and PrDOS). 3D surface hydrophobicity and charge visualization of the HLH-domain with residues positively charged in blue, negatively charged in red and hydrophobic in yellow. The Helical wheels plots of Helix 1 (aa144-161) and Helix2 (aa262-279) show hydrophobic and polar faces. C. Confocal slice of parasites expressing NCR1-GFP and second copy of either full-length NCR1 in NCR1FL parasites (top) or HLH domain truncated PfNCR1 in NCR1ΔHLH parasites (bottom). NCR1FL - or NCR1ΔHLH-mRuby3 (magenta), NCR1-GFP (green). Plots show the fluorescence signal along the parasite periphery used for PCC calculations in D. Cells are chosen to represent the range of data shown in D. Top to bottom: NCR1FL - relatively high: 0.90, average: 0.82, low: 0.66 and NCR1ΔHLH - relatively high: 0.63, average: 0.11, low: - 0.16. Protein domain diagrams on top of sub-panels introduce PfNCR1 variants expressed as second copy in the NCR1-GFP parent line: full-length NCR1 (NCR1FL) and a mutant with the HLH-domain truncated (NCR1ΔHLH). Scale bar: 5 µm. D. Pearson correlation coefficients of NCR1ΔHLH-mRuby3—NCR1-GFP (N= 33 cells) in comparison to NCR1FL-mRuby3—NCR1-GFP (N= 32 cells). Pearson correlation coefficients of EXP2-mNeonGreen—PV-mRuby3 (N= 31 cells) and EXP2-mRuby3—NCR1-GFP (N= 29 cells) parasites used to define narrow membrane contact sites for reference. Bars: median ± 95% CI. Stars: Dunnett test after ANOVA (n.s.: not significant, *: p≤0.05, **: p≤0.01, ***: p≤0.001, ****: p≤0.0001).

To characterize the structural elements of the conserved yet unresolved region, we performed in silico analysis using HeliQuest (Gautier et al. 2008). Helical wheel projections revealed amphipathicity within the α-helix residues aa144-161 and aa262-279, each with contrasting hydrophilic and hydrophobic faces (Fig. 1B). Thus, the unresolved aa142-282 region seems to have amphipathicity at both its termini, potentially capable of associating with both the hydrophobic membrane core and the aqueous PV lumen while the lysine and arginine residues may enhance the membrane affinity by association with negatively charged lipids (Drin and Antonny 2010). The flexible linker on the other hand is disordered (Fig. 1B) but enriched in negatively charged aspartic and glutamic acid repeats that can facilitate interaction with positively charged features (Bigman et al. 2022).

To assess the contact site targeting function of aa142-282 (hereafter referred to as helix-linker-helix domain, or “HLH domain”), we compared localization of mRuby3-tagged HLH domain-deleted PfNCR1 (PfNCR1ΔHLH-mRuby3) with that of a full-length version of the protein (PfNCR1FL-mRuby3). To enable colocalization experiments, the respective proteins are expressed in individual cell lines as a second copy at the well characterized *cg6* locus (Adjalley et al. 2011) in the background of endogenous GFP-tagged PfNCR1 (Garten et al. 2020). The parental cell line is referred to as NCR1-GFP^apt^, abbreviated to NCR1-GFP, and allows regulation of endogenous PfNCR1 expression via the TetR-DOZI repression system (Ganesan et al. 2016; Istvan et al. 2019), enabling assessment of function of the respective modified second copy version of PfNCR1 later on. The cell lines generated for the initial localization assessment are referred to as NCR1-GFP^apt^::NCR1ΔHLH-mRuby3, abbreviated to NCR1ΔHLH, and NCR1-GFP^apt^::NCR1FL-mRuby3, abbreviated to NCR1FL, respectively (Fig. 1C and SI Fig. S2).

Live-cell confocal microscopy showed that while PfNCR1FL-mRuby3 co-localized with PfNCR1-GFP specifically at the contact sites as expected, PfNCR1ΔHLH-mRuby3 was distributed all over the periphery of the parasite (Fig. 1C). To quantify the degree of colocalization in between endogenous GFP-labeled protein and the second mRuby3-labeled copy, the Pearson-correlation coefficient (PCC) was calculated at the periphery of the parasite, providing a colocalization scale of PCC of 1 = perfect colocalization, PPC of -1 = perfect anti-localization, PCC of 0 = random localization (SI Fig. S3). We found a PCC between PfNCR1ΔHLH-mRuby3 and PfNCR1-GFP was 0.21 [0.29, -0.02] (median [95% CI]). In contrast, PfNCR1FL-mRuby3 and PfNCR1-GFP colocalized with a PCC of 0.81 [0.85, 0.70] (Fig. 1D). To contextualize, we reproduced the PCCs of previously published cells lines (Garten et al. 2020) in our present experimental conditions: EXP2-mNeonGreen::PV-mRuby3 show a high degree of colocalization with PCC 0.82 [0.85, 0.76], while EXP2-mRuby3::NCR1-GFP shows a high degree of anti-localization with PCC -0.70 [-0.61, -0.78] (SI Fig. S4).

The colocalization of the full-length second copy and endogenous PfNCR1 is as high (PCC > 0.8) as the colocalization of EXP2 and the wider PV regions, while the localization of the HLH domain truncated PfNCR1 tends to be randomized around the parasite. Taken together, the HLH domain can be characterized as essential for PfNCR1 targeting.

### Contact site targeting requires combined contribution of the domain sub-regions

The HLH domain is 141aa in length with multiple identifiable sub-regions of different physicochemical character: helix 1 and 2 are amphipathic with a positively charged hydrophilic face, and the linker (aa186-255) in between the helices is disordered with a stretch (aa213-232) of negative charges (0 positive, 12 negative residues) (Fig. 2A).

**Fig. 2.**
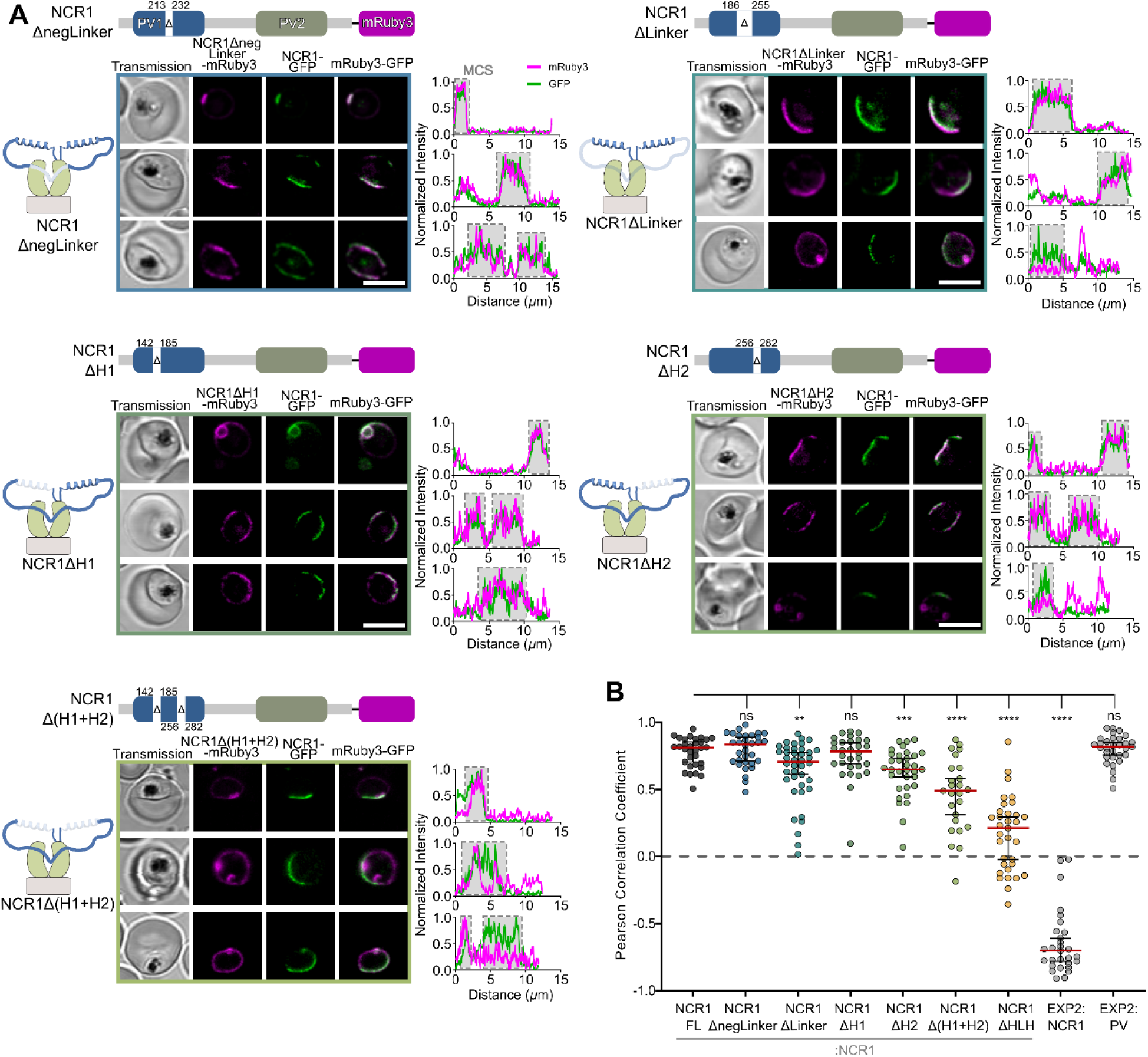
Contact site targeting requires combined contribution of the domain sub-regions. A. Confocal slice of parasites expressing NCR1-GFP and second copy of the PfNCR1 missing domain sub-regions as indicated on top of the sub panels. Plots show the fluorescence signal along the parasite periphery used for PCC calculations in B. Cells are chosen to represent the range of PCCs shown in B: Top to bottom: NCR1ΔnegLinker - relatively high: 0.98, average: 0.87, low: 0.66; NCR1ΔLinker - relatively high: 0.91, average: 0.69, low: 0.08; NCR1ΔH1 - relatively high: 0.92, average: 0.73, low: 0.61; NCR1ΔH2 - relatively high: 0.85, average: 0.68, low: 0.07; NCR1Δ(H1+H2) - relatively high: 0.80, average: 0.47, low: 0.07. Protein domain diagrams on top of sub-panels introduce PfNCR1 missing domain sub-region variants expressed as second copy in the NCR1-GFP parent line. Scale bar: 5 µm. B. Pearson correlation coefficients of NCR1ΔnegLinker-mRuby3—NCR1-GFP (N= 32 cells); NCR1ΔLinker-mRuby3—NCR1-GFP (N= 40 cells); NCR1ΔH1-mRuby3—NCR1-GFP (N= 30 cells); NCR1ΔH2-mRuby3—NCR1-GFP (N= 33 cells); NCR1Δ(H1+H2)-mRuby3—NCR1-GFP (N= 25 cells) in comparison to NCR1FL -mRuby3—NCR1-GFP (N= 32 cells). Pearson correlation coefficients of EXP2-mNeonGreen—PV-mRuby3 (N= 31 cells) and EXP2-mRuby3—NCR1-GFP (N= 29 cells) parasites shown for reference. Bars: median ± 95% CI. Stars: Dunnett test after ANOVA (n.s.: not significant, *: p≤0.05, **: p≤0.01, ***: p≤0.001, ****: p≤0.0001).

To understand the contribution of the subregions to PfNCR1 targeting, we characterized localization of PfNCR1 carrying deletions of those sub-regions. All variants were expressed in the same background as the HLH domain-truncated PfNCR1-mRuby3 (SI Fig. S2). The colocalization of each variant with respect to the endogenous PfNCR1-GFP was observed using live-cell confocal microscopy and quantified by calculating the PCC (Fig. 2B). The PCC of PfNCR1-GFP and PfNCR1ΔnegLinker-mRuby3 (aa213-232 deleted) was 0.84 [0.89, 0.71], PfNCR1ΔLinker-mRuby3 (aa186-255 deleted) was 0.70 [0.77, 0.61], PfNCR1ΔH1-mRuby3 (aa142-185 deleted) was 0.78 [0.85, 0.70], PfNCR1ΔH2-mRuby3 (aa256-282 deleted) was 0.65 [0.73, 0.60], and PfNCR1Δ(H1+H2)-mRuby3 (aa142-185 and aa256-282 deleted) was 0.49 [0.58, 0.31]. PfNCR1 with deletion of the whole disordered linker, helix 2 and both helices resulted in significant change in localization, with the deletion of the 2 helices having the largest effect of any partial deletion. The PCC of the 2x helix deletion line NCR1Δ(H1+H2) sits in between NCR1FL and NCR1ΔHLH, with the localization changing enough to separate the 95% confidence intervals of the 2x helix deletion when compared to the full-length line. It appears that each part of the HLH domain is contributing to PfNCR1 targeting. While the helices are distinguished in their contribution, the combined effect of the deletion of the sub-regions amounts to the largest change in character of PfNCR1 targeting.

### PfNCR1 mistargeting causes partial loss of cholesterol transport function

Correct localization of PfNCR1 may reasonably be expected to be important for its function in maintaining PPM cholesterol homeostasis, or it might serve a yet unknown purpose. The ‘death phenotype’ of a PfNCR1 knock-down is slow, with effects manifesting after several cycles (Istvan et al. 2019). Indeed, in our experimental conditions we could grow parasites without anhydrotetracycline (aTc), knocking down endogenous PfNCR1 without a second copy of PfNCR1, at ∼0.91x the growth rate compared to the +aTc condition (SI Fig. S5). In a functional assay, PfNCR1’s critical role in cholesterol homeostasis of the outer leaflet of the PPM has previously been demonstrated by exploiting the high affinity of saponin to cholesterol, causing leakage of cytoplasmic content (such as aldolase) if the plasma membrane contains an ‘above threshold’ amount of cholesterol for permeabilization by saponin (Istvan et al. 2019) (Fig. 3A).

**Fig. 3.**
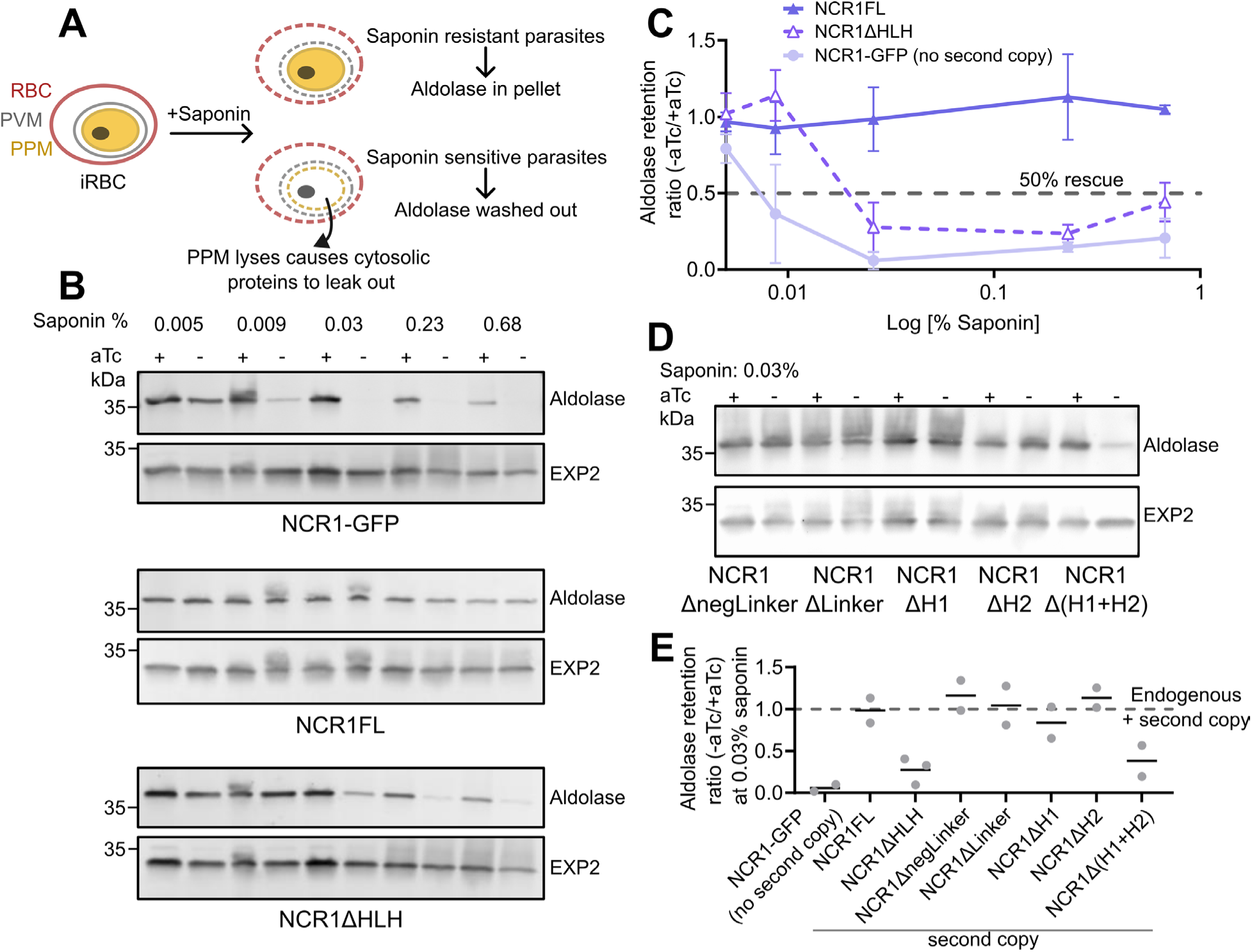
NCR1*Δ*HLH supports partial rescue of cholesterol homeostasis after knock-down of endogenous PfNCR1. A. Schematic of the saponin-sensitivity assay illustrating permeabilized membranes with and without cholesterol in the PPM. Resistant parasites: saponin forms pores on the RBC and PVM membranes, but the PPM remains intact leading to cytosolic proteins like aldolase to be retained in the parasite pellet. Sensitive parasites: saponin forms pores on the RBC, PVM and PPM causing cytosolic proteins to leak out, leading to aldolase wash-out. B. Saponin-sensitivity assay of parasites expressing NCR1 variants in the background of aTc-regulatable endogenous PfNCR1. NCR1-GFP: no second copy of PfNCR1, NCR1FL: full-length second copy of PfNCR1, NCR1ΔHLH: second copy of PfNCR1 lacking the HLH domain. PfNCR1 variants cultured with (+) or without (-) aTc were treated with increasing concentrations of saponin (0.005%, 0.009%, 0.03%, 0.23%, 0.68%) probing release of aldolase. PVM protein EXP2 is used as loading control. C. Saponin dose-response curves: Aldolase release in -aTc over +aTc parasites as a function of percentage saponin from variants NCR1-GFP, NCR1FL, and NCR1ΔHLH (Aldolase signal was first normalized to EXP2 and then the ratio of +aTc to -aTc was calculated). Data points are mean and errors are standard deviation from biological duplicate (NCR1FL, NCR1-GFP) and triplicate (NCR1ΔHLH) experiments. D. Saponin-sensitivity assay of parasites expressing deletion variants of NCR1 HLH domain sub-region under aTc regulation. NCR1 variants cultured with (+) or without (-) aTc were treated with 0.03% saponin. Antibodies as in A. E. Plot summarizing the relative aldolase levels in –aTc over +aTc parasites under mild saponin concentrations of 0.03%. Data points are mean from biological duplicate (NCR1FL, NCR1-GFP, NCR1ΔnegLinker, NCR1ΔLinker, NCR1ΔH1, NCR1ΔH2, NCR1Δ(H1+H2)) and triplicate (NCR1ΔHLH) experiments.

To test whether PfNCR1’s role in maintaining low PPM cholesterol relies on its contact site targeting ability, we performed the saponin lysis assay on parasites expressing the mislocalized PfNCR1 variant NCR1ΔHLH in the background of aTc regulatable endogenous *ncr1* expression. Absence of detectable endogenous PfNCR1-GFP was confirmed in each experiment by confocal microscopy. A saponin lysis dose – response curve was obtained from parasites grown in the presence (endogenous PfNCR1 on) and absence (endogenous PfNCR1 off) of aTc. Leakage of cytosolic aldolase was quantified using western blotting. As positive and negative controls, parasites expressing a second copy of full-length PfNCR1 (NCR1FL) or no second copy (NCR1-GFP) were used respectively (Fig. 3B, C, SI Fig. S6 A). While NCR1-GFP parasites grown in the absence of aTc were sensitive to saponin (indicated by nearly complete leakage of aldolase under mild saponin concentrations of 0.009%), the NCR1FL parasites showed resistance to aTc withdrawal even under high saponin doses (up to 0.68% saponin). In contrast, NCR1ΔHLH parasites grown in the absence of aTc showed intermediate saponin sensitivity (incomplete aldolase release at 0.03% and above) (Fig. 3C, E). These results suggest that efficient targeting of NCR1 to the contact sites is required to maintain low cholesterol levels in the PPM, and loss of contact site targeting impairs cholesterol transport function and enhances saponin susceptibility. To further determine whether milder mistargeting affects cholesterol transport function of PfNCR1, or perhaps absence of parts of the HLH-domain leads to a functional defect independent of mislocalization, parasites lines NCR1ΔnegLinker, NCR ΔLinker, NCR ΔH1, NCR1ΔH2 and NCR1Δ(H1+H2) were tested for saponin sensitivity. Treatment with 0.03% saponin, a concentration that exposes the intermediate saponin sensitivity phenotype for NCR1ΔHLH, did not result in lysis for the other truncations with exception of the 2x helix truncation, which showed saponin sensitivity similar to the HLH domain truncation (Fig. 3D, E SI Fig. S6 B).

Taken together the saponin lysis assay establishes two findings: 1) the truncated PfNCR1 is functional. Removing parts or the entire HLH domain (i.e. ∼10% of the wild type PfNCR1) results in a protein that can still functionally rescue knock-down of endogenous PfNCR1, at least partially. With this we argue that neither individual sub-regions nor the HLH domain as a whole has a critical structural role. 2) The observed intermediate saponin sensitivity of both NCR1ΔHLH and NCR1Δ(H1+H2) correlates with reduced contact site targeting and implies that correct localization of PfNCR1 is necessary for efficient maintenance of cholesterol homeostasis.

### PfNCR1 HLH domain alone targets PV-loops

To determine whether the HLH domain in whole or part is sufficient for contact site targeting, we generated parasite lines exporting the HLH domain and disordered linker sub-region to the PV. To that end, we fused the HLH-domain (aa142-282) and the disordered linker sub-region (aa186-255) at the N-terminus to a PV targeting motif derived from HSP101 as in ref (Garten et al. 2020) and tagged C-terminally with mRuby3. Expressed in the NCR1-GFP line to allow assessment of colocalization as in the previous sections, this created the cell lines PV-HLH and PV-Linker respectively. As a point of comparison, the PV-mRuby3 (mRuby3 alone fused to the HSP101 PV signal motif)::PfNCR1-GFP (cell line abbreviated to PV-mRuby3) characterized in (Garten et al. 2020) was used. As observed previously, PV-mRuby3 localized to the parasite periphery but outside the narrow contact site regions defined by PfNCR1-GFP (Fig. 4A, Fig. S7A, top panels). PV-Linker-mRuby3 was not detectable at the parasite periphery and was instead found in the parasite’s lysosomal compartment known as the food vacuole (FV) (Fig. 4A, Fig. S7A, lower panels). PV-HLH-mRuby3 fusion was retained at the parasite periphery, however it was predominantly found in structures of vesicular appearance, seemingly attached to the parasite, reminiscent of observations described as PV-loops or tubulo-vesicular network (TVN) (Charnaud et al. 2018; Wickert and Krohne 2024) but absent from the PfNCR1-GFP positive contact site regions (Fig. 4A, Fig. S7A, middle panels). Quantification of colocalization outside of the PV-loops assigned a negative correlation between PfNCR1-GFP and PV-mRuby3 to PCC = -0.57 [-0.29, -0.75], and between PfNCR1-GFP and PV-HLH-mRuby3 to PCC = -0.26 [-0.06, -0.50] (Fig. 4E). The PCC value of PV-HLH indicates localization towards wider PV regions while at the same time targeting majorly to the PV derived “loop” structures.

**Fig. 4.**
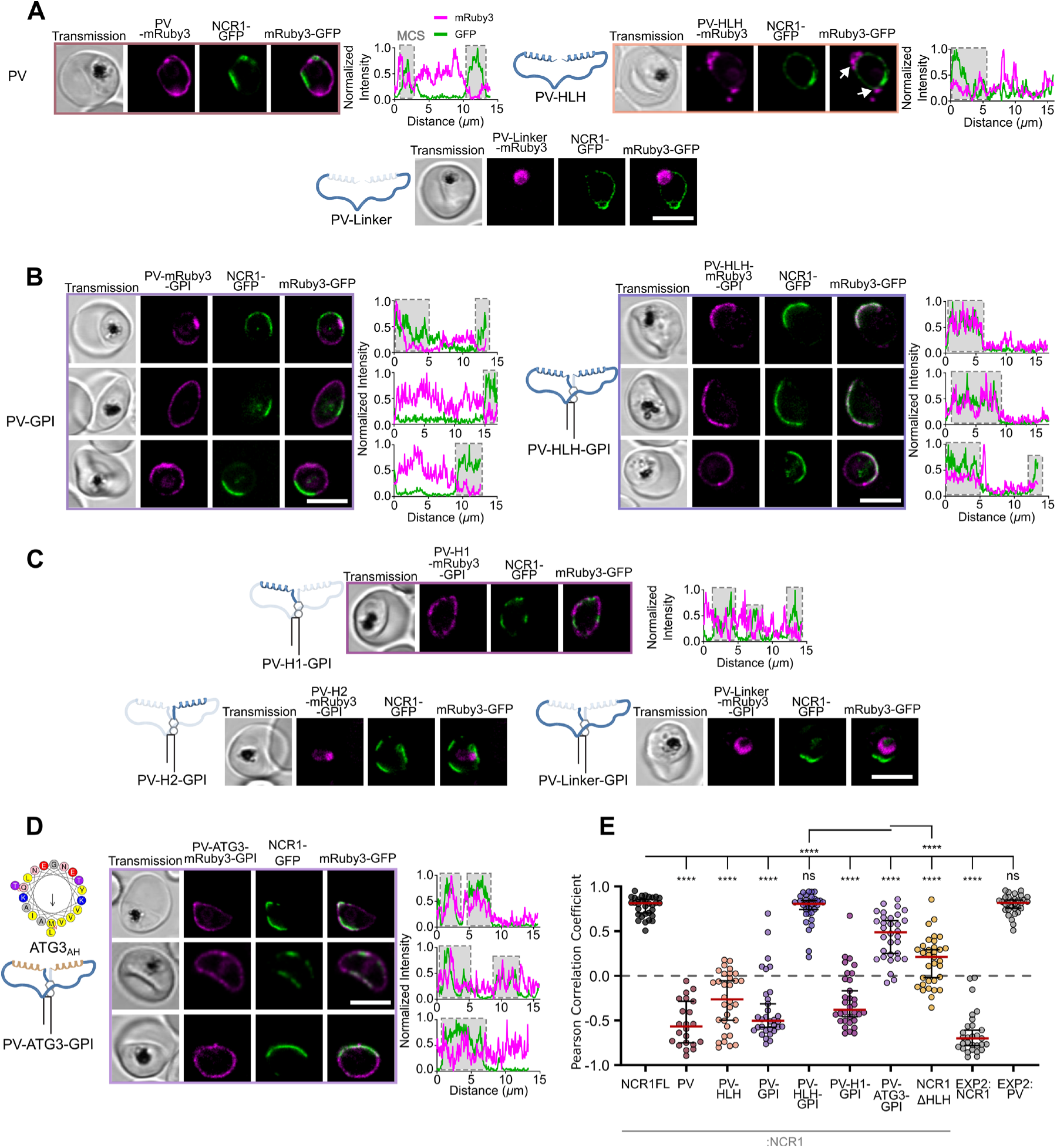
HLH-domain alone is sufficient for contact site targeting when anchored to the parasite membrane, relying on amphipathicity rather than sequence motif. A. Confocal microscopy images of parasites expressing NCR1-GFP and PV-targeted mRuby3 (PV-mRuby3), disordered linker-mRuby3 (PV-Linker-mRuby3) and the HLH-domain-mRuby3 (PV-HLH-mRuby3). Arrows highlight the localization of the HLH-domain in PV-loops. B. and C. Addition of a GPI-anchor to the tested domain variants shown in A introducing individual helices H1 and H2: PV-H1-mRuby3-GPI and PV-H2-mRuby3-GPI. C. Replacing H1 and H2 in PV-HLH-mRuby3 with ATG3_AH_ (PV-ATG3-GPI). A-D. Plots show the fluorescence signal along the parasite periphery used for E. Cells are chosen to represent the range of data shown in E, additional examples of cells in A and C can be found in SI Fig 7. Top to bottom: PV - relatively high: -0.26, average: -0.59, low: -0.78; PV-HLH - relatively high: 0.06, average: -0.44, low: -0.79; PV-GPI - relatively high: 0.19, average: -0.49, low: -0.66; PV-HLH-GPI - relatively high: 0.88, average: 0.79, low: 0.51; PV-H1-GPI - relatively high: 0.21, average: -0.35, low: -0.36; PV-ATG3-GPI - relatively high: 0.78, average: 0.47, low: -0.08. Protein domain diagrams on top of sub-panels introduce PV signal sequence tagged proteins with or without GPI expressed as second copy in the NCR1-GFP parent line. Scale bar: 5 µm. E. Pearson correlation coefficients of the cell lines with peripheral localization of the mRuby3 labeled protein calculated in between the mRuby3 and endogenous NCR1-GFP: PV-mRuby3 (N= 22 cells), PV-HLH-mRuby3 (N= 32 cells) from A; PV-mRuby3-GPI (N= 32 cells), PV-HLH-mRuby3-GPI (N= 32 cells) from B; PV-H1-mRuby3-GPI (N= 32 cells) from C; and PV-ATG3-GPI (N= 32 cells) from D; as shown as reference: For reference: colocalization with respect to EXP2: EXP2-mNeonGreen—PV-mRuby3 (N= 31 cells) and EXP2-mRuby3—NCR1-GFP (N= 29 cells). Bars: median ± 95% CI. Stars: Dunnett test after ANOVA for comparison to NCR1FL and Welch’s t test for individual comparison to ATG3 (n.s.: not significant, *: p≤0.05, **: p≤0.01, ***: p≤0.001, ****: p≤0.0001).

Observing the behavior of the HLH domain and the linker gives two valuable insights. 1) Without the helices, the linker is sequestered to the parasite FV. For a hydrophilic protein that is similar in this respect to PV-mRuby3 this is unexpected. It remains to be seen if the linker is recognized as misfolded, or the linker can serve as signal for trafficking through the PV to the FV. 2) The HLH domain alone is unable to target the narrow membrane contact site regions and is largely sequestered away from the PV in immediate contact with the parasite. This is however not an unfamiliar behavior since it has been observed for instance that an unconstrained amphipathic helix can associate with a secondary region that better satisfies its affinity profile (Lee et al. 2024).

### PfNCR1 domain anchored to the parasite membrane is sufficient for contact site targeting

The HLH domain is essential for PfNCR1 targeting, with the rest of PfNCR1 providing the potentially important spatial constraint of a membrane anchor. To test the impact of membrane constraint and geometry on protein targeting to narrow contact site regions, we constructed a fusion protein with a glycosylphosphatidylinositol (GPI) anchor for PPM attachment (Bullen et al. 2022). The anchor is followed by mRuby3 to serve both as a spacer, approximating the size of PfNCR1’s globular PV-facing domain, and also as a fluorescent tag. A version of the HLH domain (see below) and finally the N-terminal PV-targeting signal complete the protein.

Just as for PV-mRuby3, the GPI-anchored PV-mRuby3-GPI without a HLH domain is excluded from the narrow contact site regions (Fig. 4B) with a PCC of -0.50 [-0.31, -0.58]. Notably however, PV-HLH-mRuby3-GPI fusion was selectively targeted to the contact sites regions and showed strong co-localization with endogenous PfNCR1-GFP (Fig. 4B). The PCC between NCR1-GFP and PV-HLH-mRuby3-GPI signals was calculated to be 0.81 [0.84, 0.74], a degree of colocalization indistinguishable from the colocalization of endogenous PfNCR1-GFP with full-length second copy of PfNCR1FL-mRuby3 (Fig. 4B, E).

To determine if a sub-region of the HLH domain is sufficient for contact site targeting, we constructed a GPI-anchored fusion protein with either of the helices or the disordered linker in place of the HLH domain. Similar to soluble PV-Linker-mRuby3, the disordered linker (PV-Linker-mRuby3-GPI) and helix 2 (PV-H2-mRuby3-GPI) expressing GPI-anchored fusion proteins, were sequestered to the parasite FV (Fig. 4C, Fig. S7B, middle and bottom panels). However, helix 1 expressing fusion protein, PV-H1-mRuby3-GPI, was retained in the PV but mostly excluded from the narrow contact site regions (Fig. 4C, Fig. S7B, top panels). The PCC between NCR1-GFP and PV-H1-mRuby3-GPI signals was calculated to be -0.38 [-0.17, -0.46] (Fig. 4E), signifying targeting outside the narrow membrane contact site region.

The experiments restricting exploration of the PV by the tested constructs show that the HLH domain is sufficient to localize PfNCR1 when the HLH domain is anchored to the PPM. In contrast to PfNCR1 targeting, where each sub-domain deletion on its own could maintain the character of PfNCR1 localization to narrow contact sites, only the combination of HLH-domain sub-regions supports co-localization of mRuby3-GPI with endogenous PfNCR1. This argues for cooperativity between the sub-regions as a prerequisite for targeting to the narrow membrane contact sites.

### Targeting to narrow membrane contact sites is not dependent on helix sequence

While the HLH domain appears essential and sufficient for targeting a protein to the narrow membrane contact site regions of *Plasmodium falciparum*, binding to the narrow contact site region can be achieved in two ways: via binding of the HLH-domain to a protein already localized to the region, or a direct membrane interaction driven by attraction of the HLH-domain with the PVM. Direct interaction with a protein would be sequence-specific while the lipid membrane interaction would accommodate binding with a domain of matching physicochemical properties. Since amphipathic helices are candidates for membrane binding, we tested for absence of sequence-specificity of the localization by substituting the amphipathic helices of PV-HLH-mRuby3-GPI with the physicochemically similar amphipathic helix of human ATG3 (referred to as PV-ATG3_AH_-linker-ATG3_AH_-mRuby3-GPI or PV-ATG3-GPI for short) (Nishimura et al. 2023). Indeed, live cell microscopy revealed colocalization with endogenous NCR1-GFP with PCC value calculated to be 0.49 [0.61, 0.25] (Fig. 4D, E). While the narrow contact site localization is significantly different when compared to PV-HLH-mRuby3-GPI, the ATG3 amphipathic helices prevent the linker domain from sequestration in the FV and, most importantly, shift the PCC above 0, targeting PV-ATG3-GPI significantly more efficiently to the narrow membrane contact sites than PfNCR1 lacking the HLH-domain.

With the amphipathic helix of human ATG3 being able to localize the GPI anchored protein to the narrow membrane contact site regions we conclude that physicochemical properties of the protein dominate localization, with details of the required properties remaining to be uncovered.

## Discussion

The localization of PfNCR1 at narrow membrane contact site regions, characterized by ∼3-4 nm aqueous space in between membranes, with a size on the order of µm^2^, and thus accessible with a light microscope, assigns PfNCR1 a unique role amongst its homologs and allows an opportunity to define the relationship of the protein’s structure with its function and environment. Deleting the 141 amino acid (aa) long domain (aa142-282) comprised of a helix-linker-helix (HLH) arrangement leads to randomization of PfNCR1’s localization at the periphery of the parasite with a corresponding decrease in cholesterol homeostasis. Of the HLH subdomains, the helices appear to contribute most to the contact site localization. The HLH domain itself is sufficient to target a protein, here mRuby3, to the narrow contact site regions, if the movement of the protein is restricted to the vicinity of the parasite plasma membrane by a GPI anchor. In contrast, it is remarkable that GPI-anchored mRuby3 without the HLH domain is excluded from the narrow membrane contact site regions despite demonstrably fitting in between PVM and PPM, arguing for exclusion of PV proteins from narrow contact site regions that is overcome by the HLH domain. The targeting is notably not dependent on the sequence of the helices as the amphipathic helix of human ATG3 can replace the *Plasmodium* helices (see Fig. 5 for a summary of the observations).

**Fig. 5.**
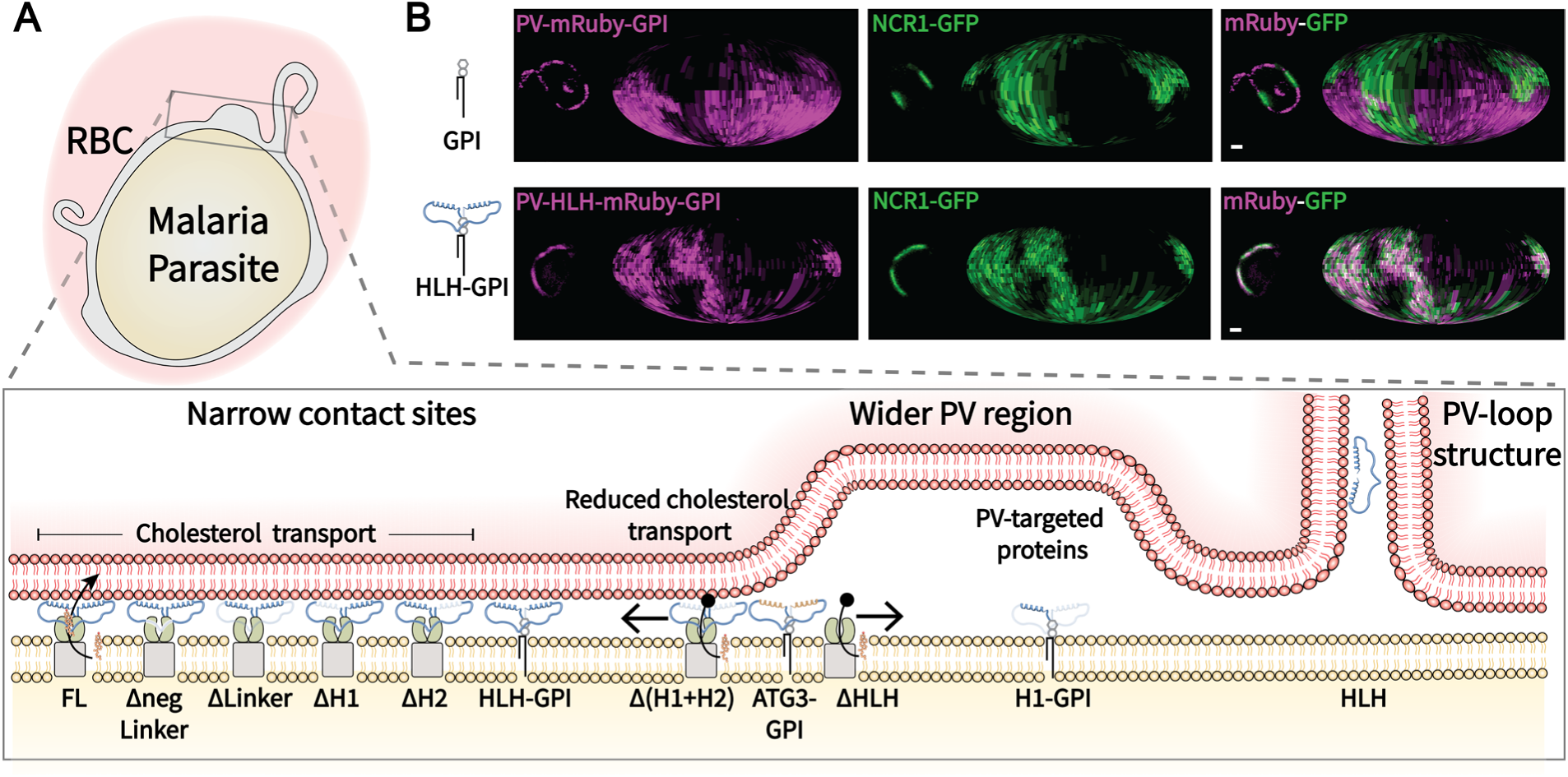
Summary of the observed targeting to regions at the *Plasmodium* – Red Blood Cell host-parasite interface. A. The work explored PfNCR1’s HLH-domain for targeting narrow membrane contact sites at the host-parasite interface of *Plasmodium falciparum* with its host-red blood cell. Removal of the HLH domain leads to randomization of PfNCR1 localization, while the HLH domain alone localized to the narrow membrane contact sites when anchored to the PPM. The localization is not sequence dependent as demonstrated by ATG3’s amphipathic helix’s ability to support targeting to the narrow contact sites. On its own the HLH will localize to PV-loops / TVN-like structures that protrude from the PV. While soluble PV-targeted proteins generally localize to regions with wider PPM-PVM distance, we found that the disordered linker domain (with and without GPI) and helix 2-GPI will be targeted to the parasite’s lysosomal ‘food vacuole’. B. Center confocal slice and Mollweide projections (Garten et al. 2020) of parasites expressing NCR1-GFP and PV-mRuby3-GPI or PV-HLH-mRuby3-GPI. Scale bar: 1 µm.

Taken together, the results suggest that the amphipathic helices directly contact a membrane in the narrow membrane contact site regions. Intuitively one can conjecture that the GPI-anchored domain is bridging the distance to the PVM exploiting the segregation of the PV into a “narrow” and “wide” zone realized by likely numerous constituents for targeting, similar to the mechanisms underlying a T-cell junction (Schmid et al. 2016). However, considering the narrowness of the contact site, matching the length of the HLH-domain, one cannot exclude that the helices bind to the PPM and sense a local property of the PPM in vicinity of PfNCR1 influenced by other proteins in the region.

Indeed, the role of amphipathic helices in sensing membrane properties, including membrane defects exposing the hydrophobic core of the membrane and lipid charge, remain the subject of current research themes (Vanni et al. 2014; Giménez-Andrés et al. 2018; Lee et al. 2024). The number of variables to explore in order to understand which properties of the amphipathic helices, in combination with the overall geometry (i.e. role of spacers and linkers, sequence and character of the linker), govern the affinity to either PVM or PPM is considerable. A more detailed investigation into the physicochemical parameters facilitating the HLH domain targeting will be insightful, potentially leading to understanding of central PVM and PPM properties, such as lipid charges or unsaturation, that drive the interactions.

Interestingly, helix 1’s aa142 – 180 are part of the Cryo-EM structure of PfNCR1 (Zhang et al. 2024). With helix 1 being resolved and its hydrophobic side facing the core of the protein, it may be that either helix 1 has a relative positioning or anchoring function for the domain relative to the rest of PfNCR1 or that helix 1 may detach to bind to the membrane leaving the stretch opposing helix 1 (∼aa292-312) exposed. The exposed stretch could in this case equally contribute to the membrane contact. It remains to be seen if this or other parts of PV1 and PV2 are involved in the protein targeting and potentially responsible for the residual localization of NCR1ΔHLH to the narrow membrane contact sites.

In this work we found two surprising targeting phenotypes. 1) The disordered linker on its own, in combination with the GPI anchor and the GPI-anchored helix 2, is targeted to the food vacuole (FV). This behavior recapitulates the trafficking of falcipain 2a and plasmepsin II (Jonsdottir et al. 2023) although in our case it remains to be demonstrated whether trafficking to the FV occurs via the PV or as part of a misfolded protein response of *Plasmodium* to exposed hydrophobic residue or aggregation. Of the two possibilities, however, a misfolded protein response seems less likely due to the HLH domain trafficking to the PV successfully. Moreover, *Plasmodium* has evolved to avoid protein aggregation, as exemplified by the protection from aggregation of asparagine-rich proteins (Muralidharan and Goldberg 2013). 2) The HLH domain without GPI anchor is targeted to PV-loops / TVN-like structures. These enigmatic, PV associated structures have been described in numerous contexts over time without assigning a consensus function (for a range of observations see for example (Lauer 1997; Przyborski et al. 2003; Sakaguchi et al. 2016; Charnaud et al. 2018)). Localization of the HLH domain is curious as it suggests that the domain’s properties cause it to be excluded from the wider PV regions, most frequently serving as final destination for exported soluble proteins and not otherwise licensed for export via PTEX (Garten et al. 2020), a behavior typically observed in liquid-liquid phase separation of proteins with disordered domains (Holehouse and Kragelund 2024). Targeting to these unexpected locations can be exploited to gain understanding of other principles guiding proteins to their target within the PV.

The saponin lysis assay probes the state of cholesterol homeostasis (Istvan et al. 2019) by exploiting the ability of saponin to disturb and porate membranes (Baumann et al. 2000). This allowed us to positively assign significance to the membrane contact for PfNCR1 function, giving critical insight into the mechanisms targeted by several promising anti-malarials (Istvan et al. 2019) and also significant for a potential uptake pathway of anti-malarials (Fraser et al. 2024). As saponin binds cholesterol directly but is not membrane permeable, one may conjecture that the saponin lysis assay probes the availability of cholesterol in the outer leaflet of the PPM and it cannot distinguish how cholesterol is being depleted by PfNCR1: cholesterol can either be transferred from the PPM to the PVM or flipped to the inner PPM leaflet. Interestingly, large modifications of the extracellular domain of the mouse hPTCH1 homolog, mPTCH1, leave the cholesterol transport function of the protein intact (Briscoe et al. 2001; Kinnebrew et al. 2021). With the absence of the HLH domain impacting localization and function, it is established that the extracellular domains of PfNCR1 are necessary for cholesterol homeostasis, however it remains unknown if the extracellular domains have a regulatory role as it is seen with hPTCH1 or have a cholesterol trafficking function. With the now uncovered targeting mechanism of PfNCR1, it is feasible to ask if and under which condition other members of the family, e.g. hNPC1 or hPTCH1, can rescue *Plasmodium* cholesterol homeostasis and gain insight into the mechanistic role of membrane contact. Conversely, rescue experiments using PfNCR1 homologs also have the potential to uncover the general mechanisms underlying the cholesterol transport function of the protein family and their function in cholesterol homeostasis of humans.

Lastly but perhaps most significantly, the engineered GPI-anchored HLH-domain allows targeting of genetically encoded probes to the narrow membrane contact site regions. In principle this ability enables exploration of this only recently defined region at the host-parasite interface, using proximity biotinylation or similar strategies. Since the narrow membrane contact sites cover about half of the *Plasmodium* Host-Parasite Interface, the targeting mechanism of PfNCR1 is key to revealing the functions realized in this major yet to be characterized region.

## Materials and Methods

### Parasite Culture

*Plasmodium falciparum* was cultured in T25 culture flasks (Avantor, PA) in RPMI 1640 supplemented with 25 mM HEPES, 0.1 mM hypoxanthine, 25 μg/ml gentamicin, 0.5% Albumax II (Gibco, MD), and 4.5 mg/ml glucose (Millipore Sigma, St. Louis, MO) at 37 °C in a 5% CO_2_, 5% O_2_ atmosphere at 2.5-5 % hematocrit. Red blood cells (RBC) were obtained from anonymous healthy donors from the Stanford Blood Center (Stanford, CA) (IRB exempt) and washed with RPMI 1640 three times to remove serum and other blood cells. Parasite lines were maintained under selection drug pressure in culture.

### Genetic modification of P. falciparum

The plasmid for the expression of aTc-regulated endogenously-tagged NCR1-GFP was generated by modification of the pMG75-NCR1 plasmid (Istvan et al. 2019). GFP was amplified by PCR from Wt-NCR1 plasmid with primers GFP-1 and GFP-2 and inserted into the AatII restriction site. The construct was linearized with AflII, co-transfected into NF54:attB parasites (Adjalley et al. 2011) with pAIO-PfNCR1-gRNA1 and selected with blasticidin as described. A clonal parasite population was obtained with limiting dilution.

For the overexpression constructs, mRuby-3XFlag was isolated using a NheI and EagI restriction digest from a plasmid containing PMV-Ruby-3xFlag (Polino et al. 2023) and cloned into the plasmid EOE-2xattP (Mukherjee et al. 2022). This base vector was digested with XhoI and NheI and the full-length sequence of the StAR gene was cloned using Infusion cloning creating plasmid StAR-pyEOE-attP-mRuby3-3xFlag. StAR was amplified from gDNA using primers StAR-1 and Star-2. The StAR-pyEOE-attP-mRuby3-3xFlag construct was modified to generate plasmids with NCR1 variants.

For generation of NCR1FL-mRuby3-3XFlag-pyEOE-attP plasmid in the NCR1-GFP: NF54:attB parasite background, *ncr1* coding sequence without introns was PCR amplified from NF54:attB derived complementary DNA with primers P1 and P4 and inserted between XhoI and NheI in the plasmid StAR-pyEOE-attP-mRuby3-3xFlag. Using the same strategy, we generated the truncated variant NCR1ΔHLH-mRuby3-3XFlag-pyEOE-attP plasmid by amplifying the *ncr1* coding sequence from amino acid positions 1-141 and 283-1470 using primers P1 and P2, and P3 and P4 respectively. To generate the truncated variant NCR1ΔpLinker-mRuby3-3XFlag-pyEOE-attP plasmid by amplifying the *ncr1* coding sequence from amino acid positions 1-212 and 233-1470 using primers P1 and P11, and P12 and P4 respectively. To generate the truncated variant NCR1ΔLinker-mRuby3-3XFlag-pyEOE-attP plasmid, we amplified the *ncr1* coding sequence from positions 1-185 and 256-1470 using primers P1 and P9, and P10 and P4 respectively. To generate the truncated variant NCR1ΔH1-mRuby3-3XFlag-pyEOE-attP plasmid, we amplified the *ncr1* coding sequence from positions 1-141 and 186-1470 using primers P1 and P34, and P35 and P4 respectively. To generate the truncated variant NCR1ΔH2-mRuby3-3XFlag-pyEOE-attP plasmid, we amplified the *ncr1* coding sequence from positions 1-255 and 283-1470 using primers P1 and P36 and P37 and P4 respectively. To generate the truncated variant NCR1Δ(H1+H2)-mRuby3-3XFlag-pyEOE-attP plasmid, we amplified the *ncr1* coding sequence from positions 1-141, 186-255, and 283-1470 using primers P1 and P34, P35 and P36, and P37 and P4 respectively. The EXP2-mNeonGreen::PV-mRuby3 and EXP2-mRuby3::NCR1-GFP parasite lines used as controls for the PCC plots were the same as in (Garten et al. 2020).

For expression of PV-targeted NCR1 variants we either utilized signal peptide from *hsp101* coding sequence from positions 1-26 (MTRRYLKYYIFVTLLFFVQVINNVLC), or *p113* coding sequence from positions 1-22 (MKIPFFILHILLLQFLLCLIRC). To generate the variants PV-Linker-mRuby3-3XFlag-pyEOE-attP and PV-HLH-mRuby3-3XFlag-pyEOE-attP in the NCR1-GFP: NF54:attB parasite background, we amplified the *ncr1* coding sequence from positions 186-255 and 142-282 using primers P18 and P19 and P7 and P8, respectively. Hsp101 signal peptide was attached to the N-terminus of these sequences by PCR amplifying the *hsp101* coding sequence from NF54:attB derived complementary DNA with primers P5 and P17 (for PV-Linker-mRuby3) and P5 and P6 (for PV-HLH-mRuby3) and assembled along with variant NCR1 peptide sequences into the plasmid StAR-pyEOE-attP-mRuby3-3xFlag between XhoI and NheI sites. The PV-mRuby3:NCR1-GFP parasite line used was the same as in (Garten et al. 2020).

For expression of NCR1 variants tagged with GPI at the C-terminus, we utilized the *p113* coding sequence from positions 931-969 (SEEDTTPNETNKTDNGSSFFFAMSNALLVILLLLFIE FL). To generate PV-Linker-mRuby3-GPI-pyEOE-attP plasmid in the NCR1-GFP: NF54:attB parasite background, we amplified the signal peptide and GPI-tag from the *p113* coding sequence from NF54^attB^ derived complementary DNA with primers P59 and P60 and P46 and P47, respectively, and fused with the Linker-mRuby3 peptide by amplification from the PV-Linker-mRuby3-pyEOE-attP-3xFlag plasmid using primers P44 and P45. NcoI and NheI sites were introduced at the N- and C-terminal of the Linker region in the PV-Linker-mRuby3-GPI plasmid, which was replaced with the *HLH* coding sequence, amplified using the primers P48 and P49, to generate PV-HLH-mRuby3-GPI-pyEOE-attP plasmid. For generation of PV-mRuby3-GPI-pyEOE-attP plasmid, signal peptide from the *p113* coding sequence was PCR amplified from NF54:attB derived complementary DNA with primers P59 and P66 and inserted between XhoI and NheI in the plasmid PV-Linker-mRuby3-GPI-pyEOE-attP. The plasmids PV-H1-mRuby3-GPI-pyEOE-attP, PV-H2-mRuby3-GPI-pyEOE-attP, and PV-ATG3_AH_-linker-ATG3_AH_-mRuby3-GPI-pyEOE-attP were prepared by GenScript through synthesis of the H1, H2 and ATG3_AH_-linker-ATG3_AH_ peptides and their insertion between the p113 signal sequence at the N-terminus and mRuby3 at the C-terminus of the PV-HLH-mRuby3-GPI-pyEOE-attP plasmid. For each construct, the sequences were stitched together using HiFi-Gibson assembly cloning kit (New England Biolabs). The sequences synthesized by GenScript are listed in Table 2.

To facilitate integration of the NCR1 variants into the benign *cg6* locus of NF54:attB parasites expressing endogenously tagged NCR1-GFP, we applied the serine integrase-mediated recombination of attB × attP. Integration of plasmids into the attB site was done as previously established (Adjalley et al. 2011). The resulting plasmids, which also contain the *attP* sequence, were co-transfected with the integrase plasmid pINT into the NCR1-GFP: NF54:attB parasite line and selected with 10 nM WR99210 (MilliporeSigma), which was applied into the culture media 48 hours post-transfection. Clones were derived for each resulting parasite line by limiting dilution. The parent NCR1-GFP parasite line expressing the TetR/DOZI aptamer system (i.e. NCR1-GFP^apt^) strains were cultured with 500 nM anhydrotetracycline (aTc). All primers are listed in Supplementary table 1. All constructed plasmids were confirmed by sequencing and integrants were verified by PCR (SI Fig 2A).

### Saponin hypersensitivity assay

To monitor sensitivity to saponin (Saponin Quillaja sp., Sigma, S4521, lot SLBV8115), parasites were cultured in the presence or absence of 500 nM aTc for 72 h. Late-trophozoite stage parasites with 5% parasitemia and 2% hematocrit were pelleted (3 min x 800 g). Pellets were suspended in 10X volume of either increasing concentrations (0.005%, 0.009%, 0.03%, 0.23%, 0.68%) of room temperature saponin or a single concentration (0.03%) for 5 minutes. The released parasites were collected by centrifugation (3 min x 2200 g) and washed five times in cold PBS. Parasites were either proceeded for western blotting or stored at –80°C until use.

### Western Blotting

For western blots monitoring saponin hypersensitivity assay, the parasite pellets were resuspended in RIPA lysis buffer (25 mM Tris (pH 7.6), 150 mM NaCl, 1% NP-50, 0.1% SDS, 1% Sodium Deoxycholate) containing 0.1% CHAPS and protease inhibitor cocktail (Roche; Cat 5892970001) and sonicated briefly. Soluble proteins after centrifugation (30 min, 17 k g) were added to sample buffer, briefly heated at 98°C and loaded onto a 4-15% gradient TGX gel (Biorad). Proteins on the gel were transferred to a PVDF membrane (Biorad) with 20% methanol. Membranes were blocked with 5% BSA and probed with primary antibodies overnight at 4°C. Primary antibodies used were rabbit α-aldolase (Abcam; Cat# AB207494) at 1:5000 and mouse α-EXP2 antibodies (The European Malaria Reagent Repository) at 1:5000.

For blots detecting expression of PfNCR1 and its variants, parasite pellets were first resuspended in MQ water containing the protease inhibitor cocktail and freeze/thawed 3 times with liquid nitrogen/42°C dry bath. Pellets containing the membrane bound proteins obtained by centrifugation (30 min,17k g) were resuspended in RIPA lysis buffer containing 0.1% CHAPS and protease inhibitor cocktail (Roche), briefly sonicated and incubated overnight at 4°C with shaking. The samples were passed through syringe 10 times, incubated at 42°C for an hour and centrifuged (30 min, 17k g) to obtain the proteins in the soluble lysate. Soluble proteins were added to sample buffer, briefly warmed at 42°C and loaded onto 4-15% gradient TGX gel (Bio-Rad). Proteins on the gel were transferred to a PVDF membrane (Bio-Rad) with 10% methanol and 0.01% SDS overnight at 4°C. Membranes were blocked with 5% BSA and probed with primary antibodies overnight at 4°C. Primary antibodies used were mouse α-Flag (MilliporeSigma; Cat# F1804) at 1:500 and mouse α-GFP antibodies (Living colors A. v.; Cat# 632381) at 1:1000.

Secondary antibodies used were goat-α-Mouse (800CW; P/N: 926-32210) and goat-α-Rabbit (680RD; P/N: 926-68071) IR-Dyes at 1:10,000 from Licor. The iBright™ FL1500 Imaging System (Invitrogen) was used to image the fluorescent Western blots.

### Structure analysis and figure assembly

Alphafold2 structure prediction (Jumper et al. 2021) of PfNCR1 (PfNCR1_AF-Q8I266-F1-model_v4) was downloaded from https://alphafold.ebi.ac.uk/entry/Q8I266 on 12/08/2022. Chimera (Pettersen et al. 2004) was used for visualization of the PfNCR1 structure prediction. PyMOL 2.5.4 (Schödinger) and Yrb-script (Hagemans et al. 2015) were used for surface hydrophobicity and charge visualization. Disordedness of the HLH-domain was calculated using PONDR VL-XT (Romero et al. 2001) and PrDOS (Ishida and Kinoshita 2007) prediction servers. Clustal omega (Sievers et al. 2011) was used to generate multiple sequence alignments for the PfNCR1 homologs. Figures 1-4 were assembled using Inkscape (1.3.2 (091e20e, 2023-11-25, custom)) and figure 5 was prepared using Adobe Illustrator (Adobe Inc. (2025), Version 30.0).

### Microscopy

Confocal microscopy was performed on an Abberior infinity line microscope (Schloetel et al. 2019). For mNeonGreen and GFP the fluorophore was excited with a 485 nm pulsed laser. For mRuby3 the fluorophore was excited with a 561 nm pulsed laser. Parasites were not synchronized for observation but chosen to be imaged when they reached later trophozoite stage (hemozoin was clearly visible but the parasite was not segmented and the PV domains become optically resolvable). Mollweide projections in Fig. 5 were prepared as in (Garten et al. 2020).

### Image analysis

To calculate the Pearson correlation coefficient images acquired at the center/equator of the parasite were used. This allowed analyzing fluorescence consistently across cell lines without issues caused by bleaching when taking volume images as needed for the Mollweide projections in (Garten et al. 2020). Signal at the periphery of the parasite was pre-selected by manually tracing the outline of the parasite in Fiji (“Fiji Is Just ImageJ” version 1.54f) and applying the Edit > Selection > Straighten with a 20 pixel line width to include background pixels in the data, resulting in a straightened images of the parasite periphery 20 pixels wide with the length of the periphery. The analysis script records the 3 highest fluorescence intensities along the width for each position along the periphery. The median of the 3 highest intensities was recorded as membrane fluorescence intensity. As the membrane was segmented to obtain the membrane fluorescence intensity, the intensity was used without further thresholding for calculation of the Pearson Correlation Coefficient. The data was plotted, and the error was calculated with GraphPad Prism (version 10.6.1). Statistical significance compared to the reference strain was calculated with a Dunnett test after one-way ANOVA (Brown-Forsythe and Welch ANOVA tests). For pair wise comparison Welch’s t test was used.

## Supporting information

source data underlying histograms, plots, and averages with errors

## Acknowledgements

We thank all members of the Garten lab and the Apicomplexan supergroup (labs of Elizabeth Egan, Prasanna Jagannathan, John Boothroyd) for constant feedback throughout the project; Ellen Yeh and Stephanie Kabeche for sharing TC space and resources while the lab was set up; Josh Beck and his lab for help and training to engineer parasites. We thank Maia Kinnebrew for insightful discussions about the mechanisms of human PTCH1 cholesterol transport. A.R. was partially supported by funding from the Stanford Bio-X Interdisciplinary Seed Grants Program (IIP) [R12] to M.G. Research reported in this publication was supported by National Institute of General Medical Sciences (NIGMS) of the National Institutes of Health under award number R35GM160156, partially supporting consumables and salaries to M.G. and A.R. Work by DEG and ESI was supported by NIAID grant R01 AI92669. The content is solely the responsibility of the authors and does not necessarily represent the official views of the National Institutes of Health.

## Data and resource availability

Data supporting the findings of this manuscript and cell lines are available from the corresponding authors upon reasonable request. The source data underlying histograms, plots, and averages with errors are provided as a Source Data file. Source data points are provided with this paper. An example image is provided with the scripts (see code availability section).

## Code availability

MATLAB (Mathworks) scripts as described in the methods are available at https://github.com/gartenm/line_coloc/.

## Author contribution

M.G. and A.R. conceived the project and designed experiments. E.I. and D.G. designed and engineered the NCR1-GFP^apt^ parent cell line. A.R. engineered all other cell lines and performed all experiments. M.G. designed the analysis pipeline, A.R. quantified all data. M.G. and A.R. wrote the manuscript. All authors discussed the results and edited the manuscript.

## Supplementary Information

**Supplementary Table 1.**
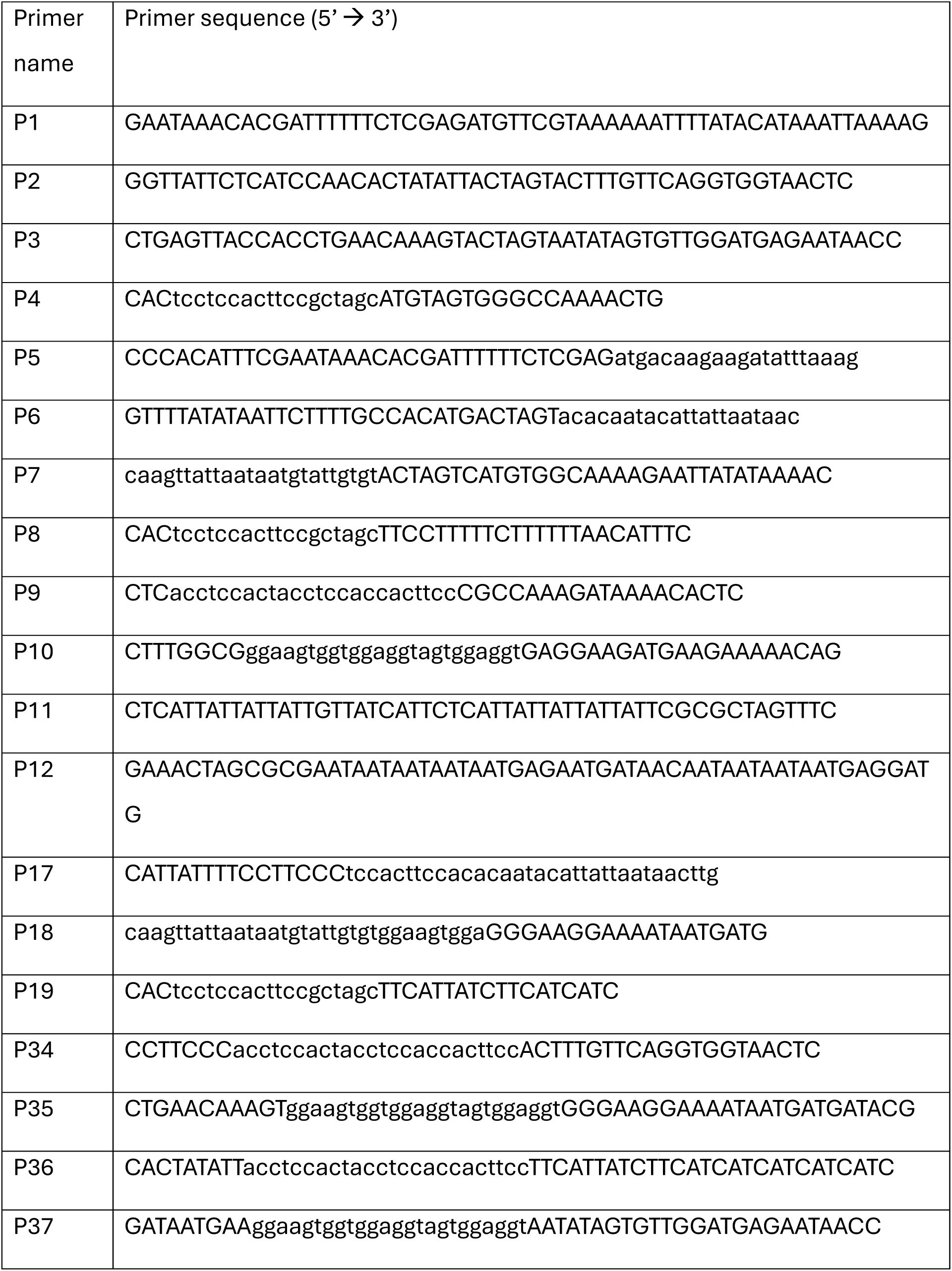
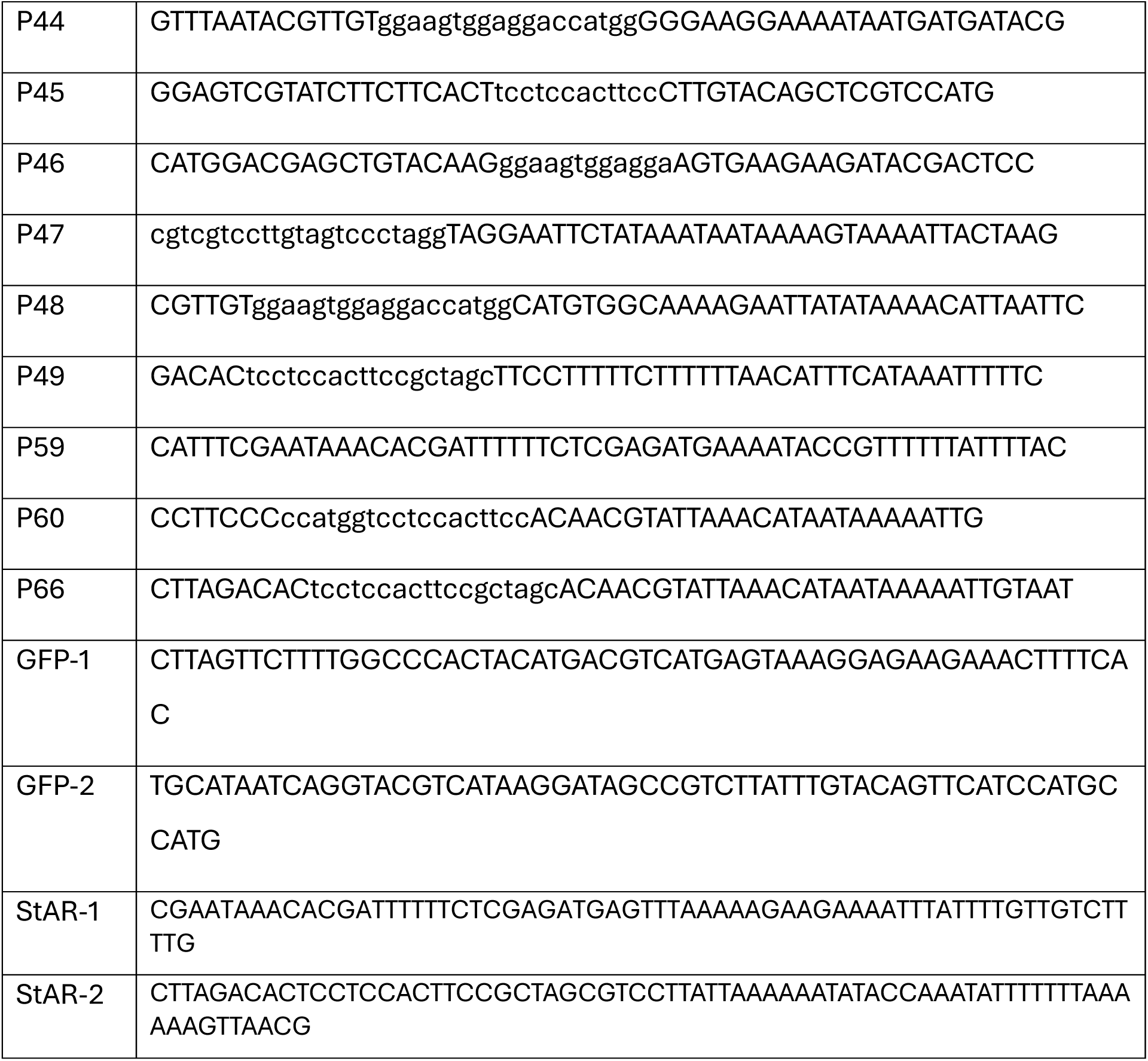

**Supplementary Table 2.**
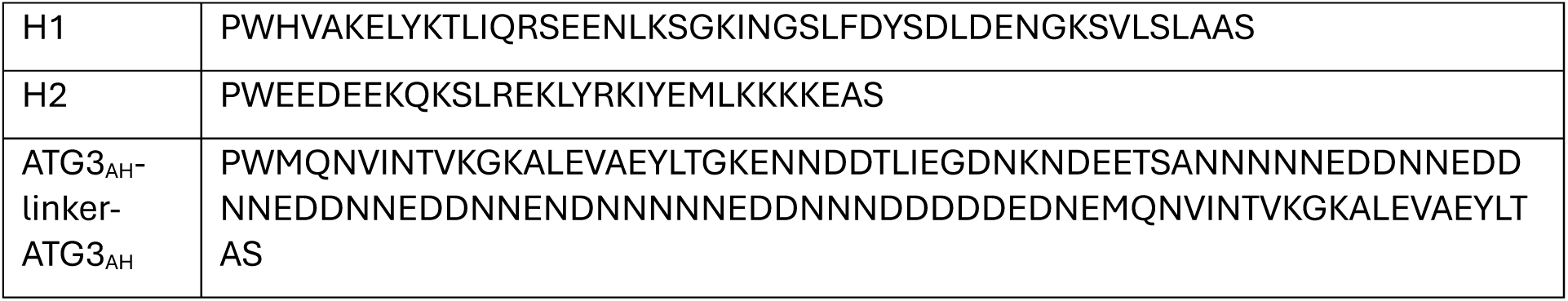

## Supplementary Figures

**Fig. S1:**
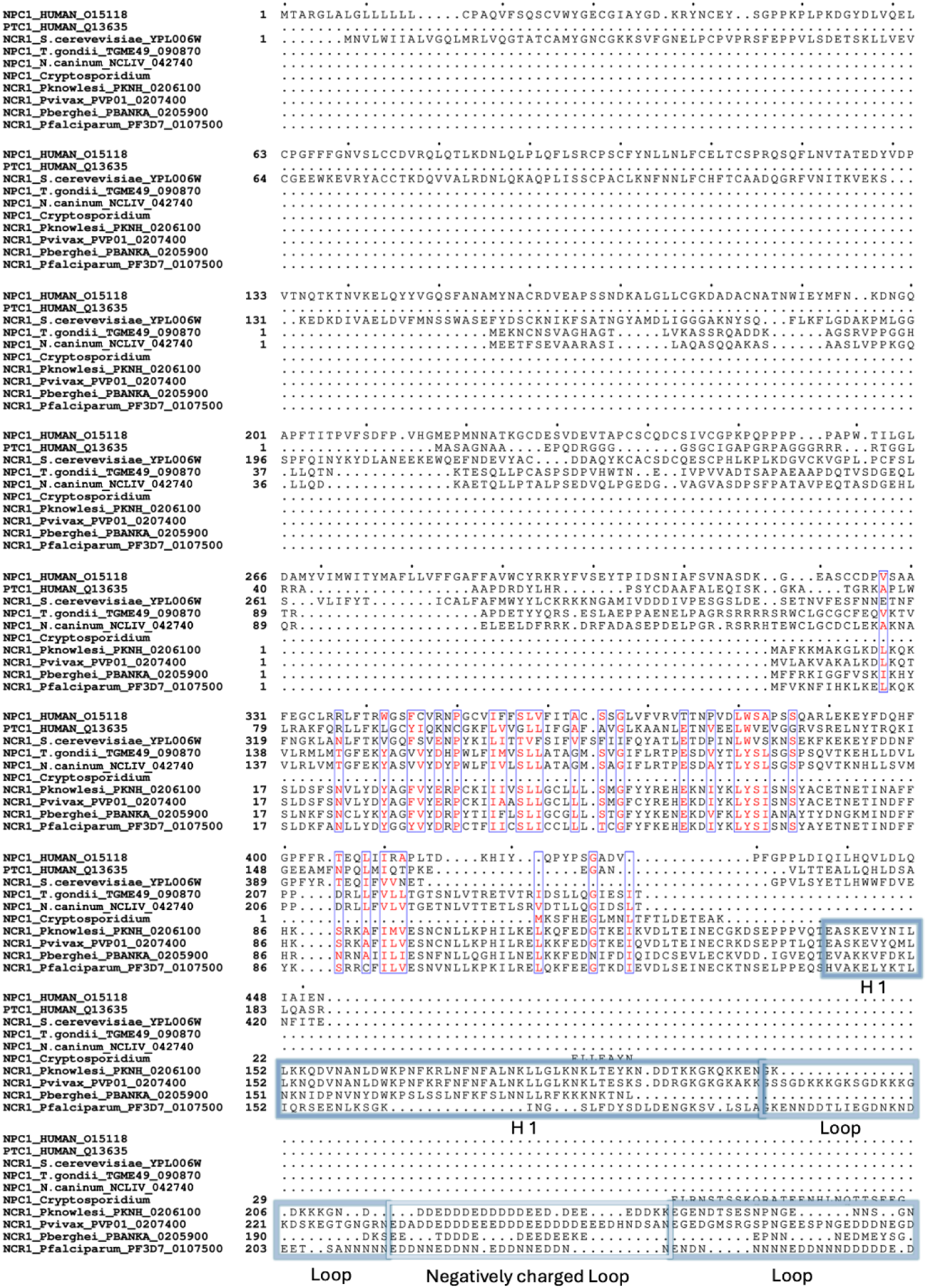
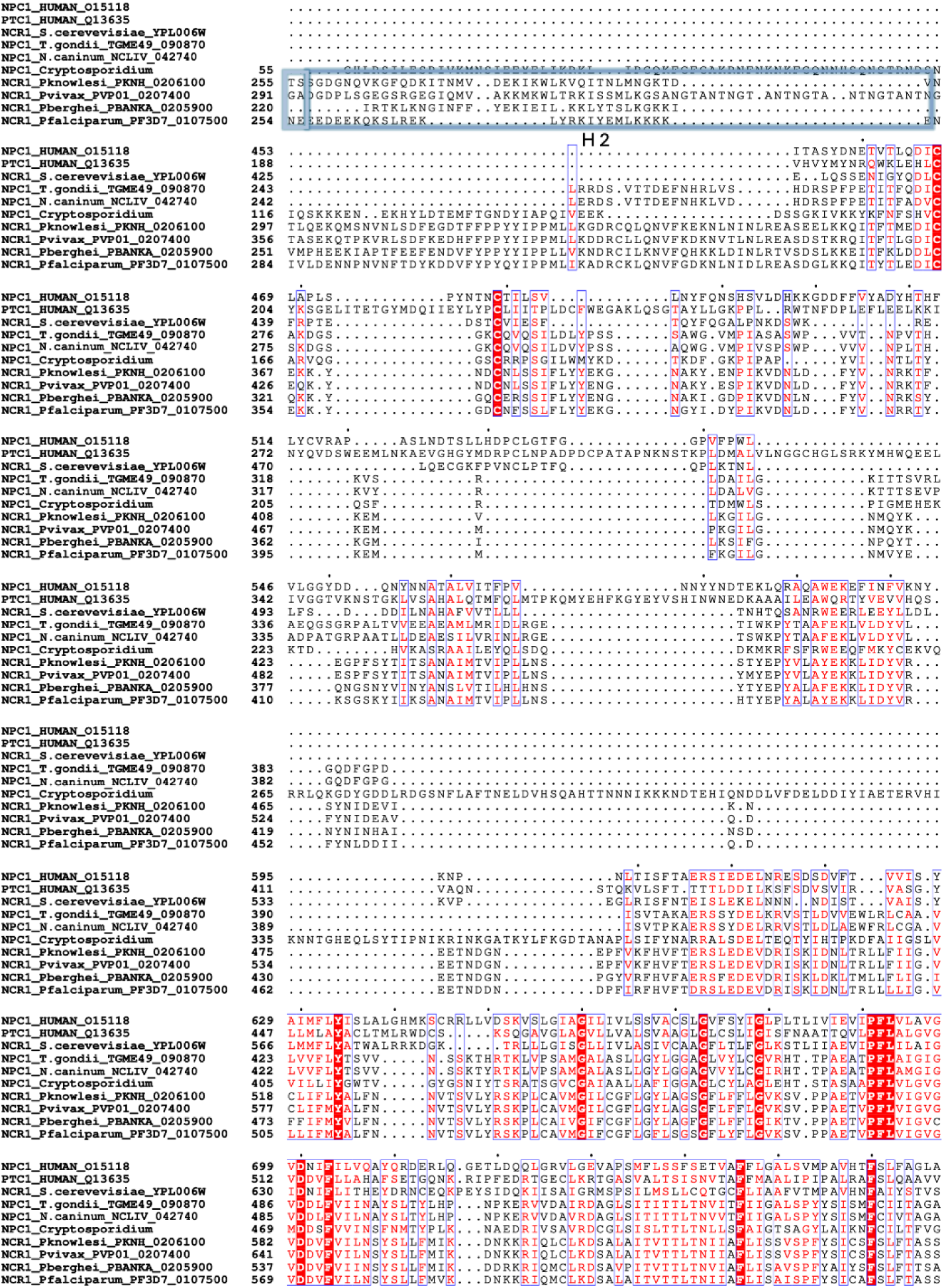
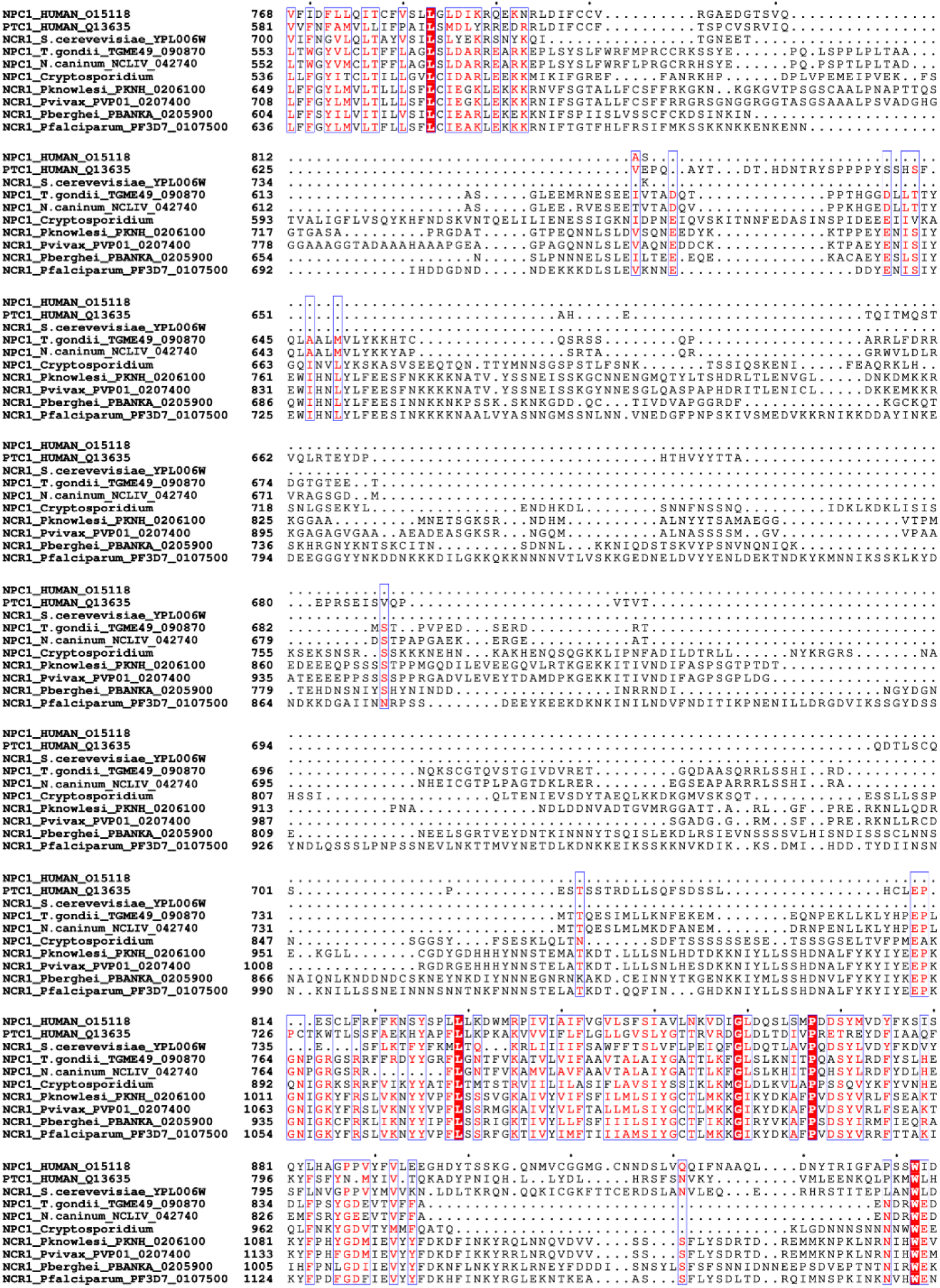
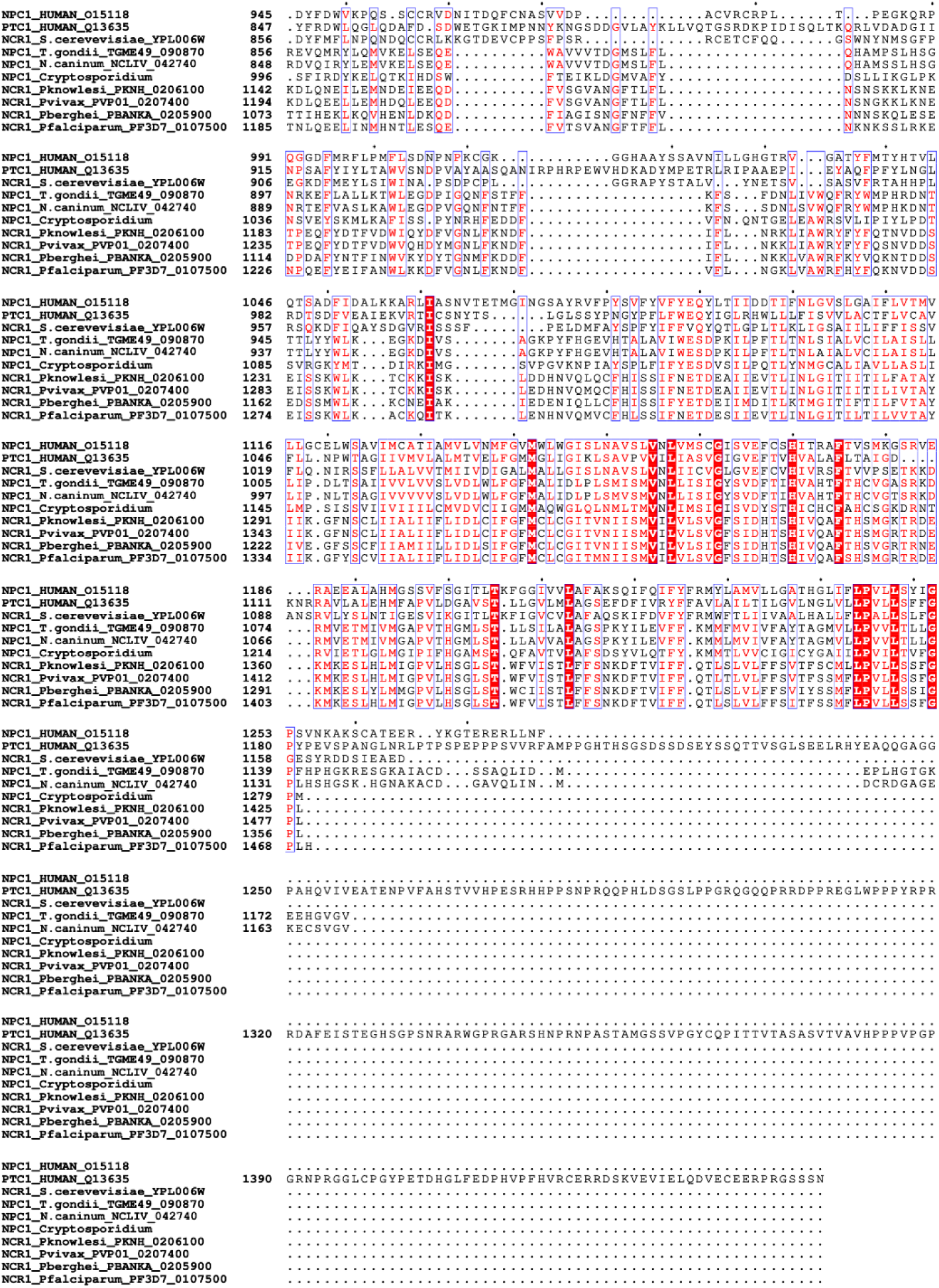
The aa142-282 region in PfNCR1 consisting of amphipathic *α*-helices bridged by a disordered linker is unique to *Plasmodium* species. Clustal-O alignment of PfNCR1 sequence with homologs from other eukaryotes. The HLH-domain is absent in homologs from human, yeast and related parasites but is conserved across *Plasmodium* species, with presence of the negatively charged aspartic and glutamic acid repeats. Regions corresponding to Helix1, Helix2, disordered linker and negatively charged repeats in the linker are boxed in shades of blue.

**Fig. S2.**
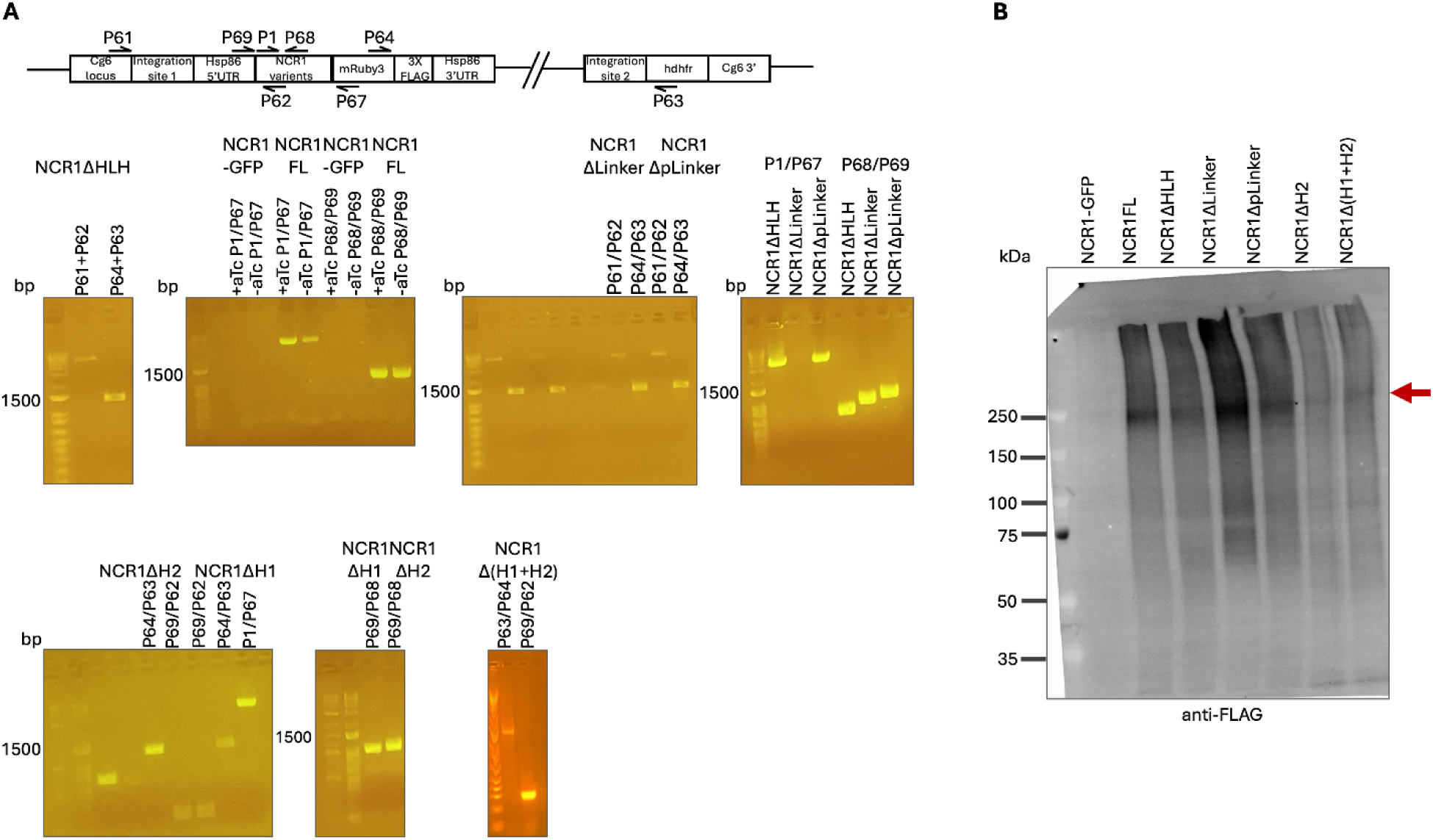
Verification of expression of second copy NCR1 variants in NCR1-GFP^apt^ parasites. A. Schematic representation of the attB × attP recombination leading to integration of ncr1 variant genes at the cg6-attB locus. PCR reactions using p61 + p62, p63 + p64, p61+p67, and p68+p69 were performed to confirm integration of the genes at the cg6-attB locus. Sequences of the primers for PCR screening are as follows: P1: 5’ GAATAAACACGATTTTTTCTCGAGATGTTCGTAAAAAATTTTATACATAAATTAAAAG, P61: 5′-GAAAATATTATTACAAAGGGTGAGG, P62: 5′-GTCGTTTATGGTTTCATTCGTTTC, P63: 5′-CTCTTCTACTCTTTCGAATTC, P64: 5′-GATGATGTATCCAGCAGATG, P67: 5′-GTACGGCTGCCATACATGAACGAC, P68: 5′-CTAAATTATCAACTTTTATAGGATAATCT ATATATCCATTTC, P69: 5′-CTTCCCACATTTCGAATA AACAC. B. Western blot analysis of the parent NCR1-GFP and second copy variants of NCR1FL, NCR1ΔHLH and HLH sub-region deletion parasite lines. Arrow indicates prominent band at ∼200 kDa in the second copy variants and no band in the parent NCR1-GFP line. The expected molecular weight of NCR1FL is 200.4 kDa while that of deltaHLH, with the largest amino acid truncation, is 183.9 kDa. When compared to NCR1FL, a difference of ∼16 kDa for NCR1ΔHLH and an even smaller difference in size for the sub-domain truncations remain unresolved, and they appear to co-migrate potentially due to limited resolution around ∼250 kDa range.

**Fig. S3.**
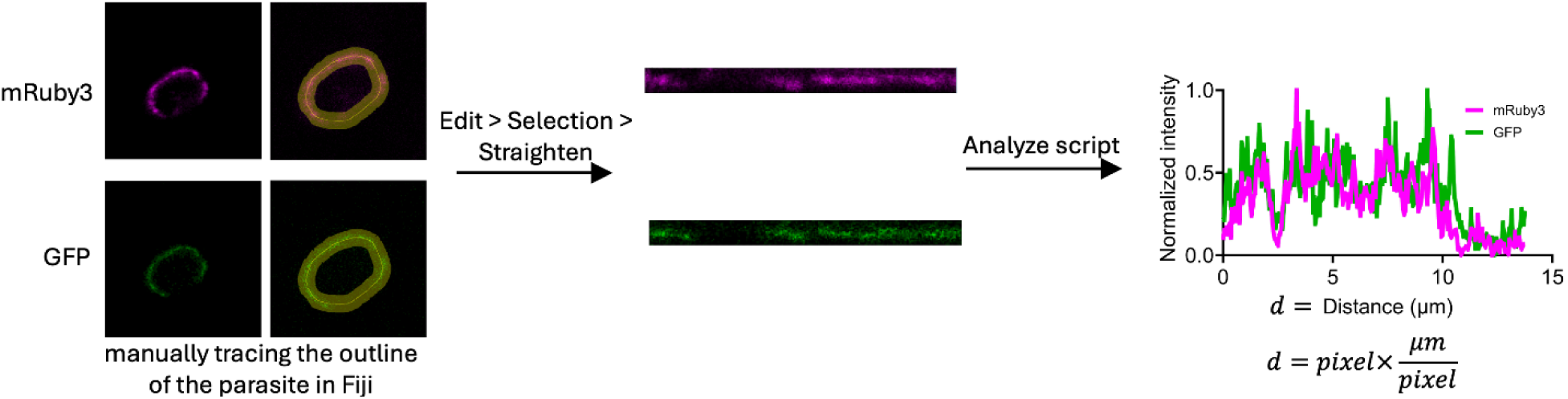
Schematic of periphery unwinding for fluorescent intensity analysis. A segmented region of interest at the parasite periphery is converted into a linearized contour of length dd=pixels* (microns/pixels), enabling position-dependent plotting of GFP and mRuby3 fluorescence intensities along the membrane.

**Fig. S4.**
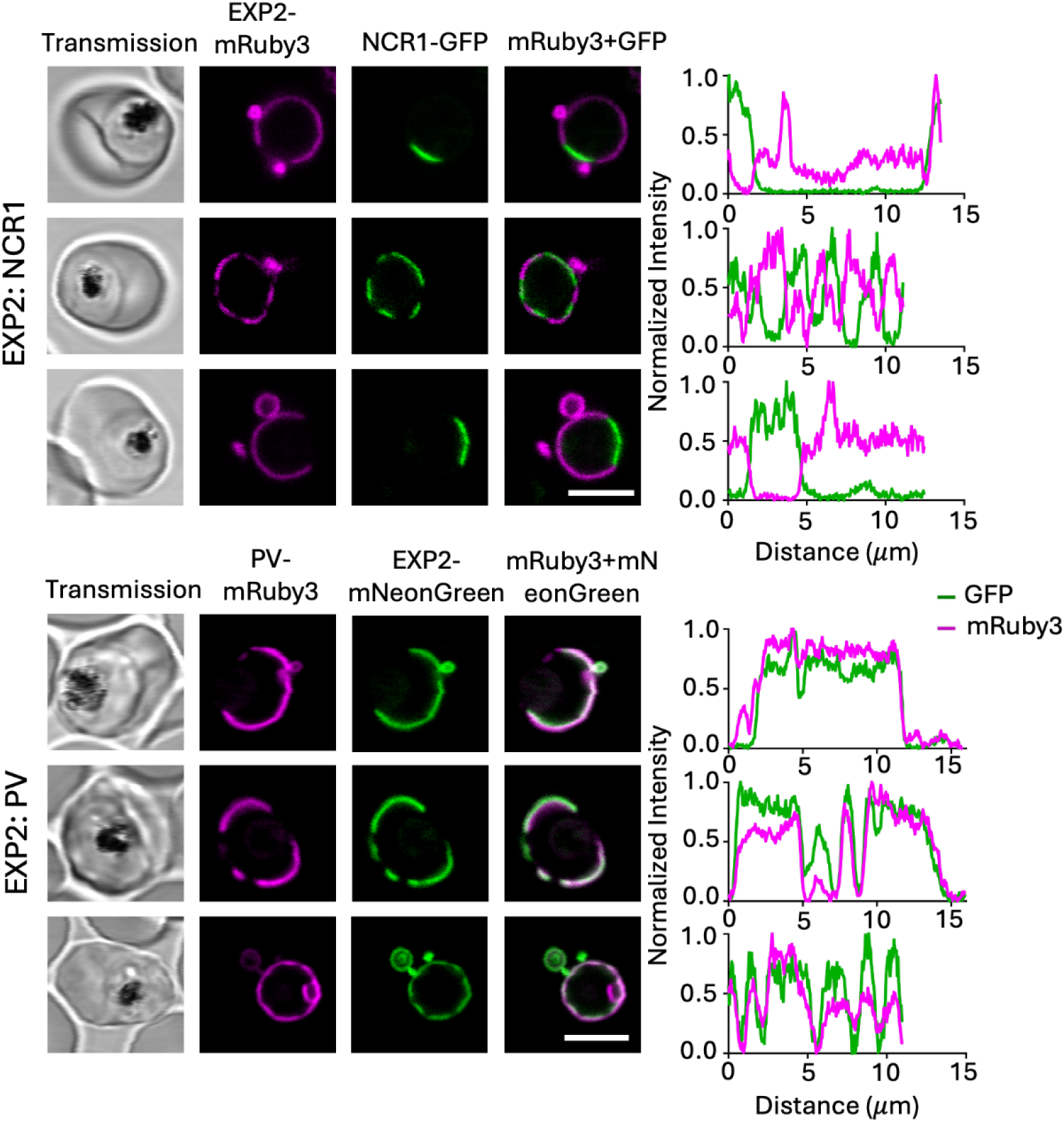
Confocal images for reproduction of PCCs of previously published cells lines (Garten et al. 2020) for our current experimental setup conditions. Confocal slice of parasites expressing NCR1-GFP::EXP2-mRuby3 (top) and EXP2-mNeongreen::PV-mRuby3 (bottom). EXP2- or PV-mRuby3 (magenta), NCR1-GFP or EXP2-mNeongreen (green). Colocalization analyzed by plotting the GFP/mNeongreen and mRuby3 fluorescence intensities measured across the parasite periphery. Scale bar: 5 µm.

**Fig. S5.**
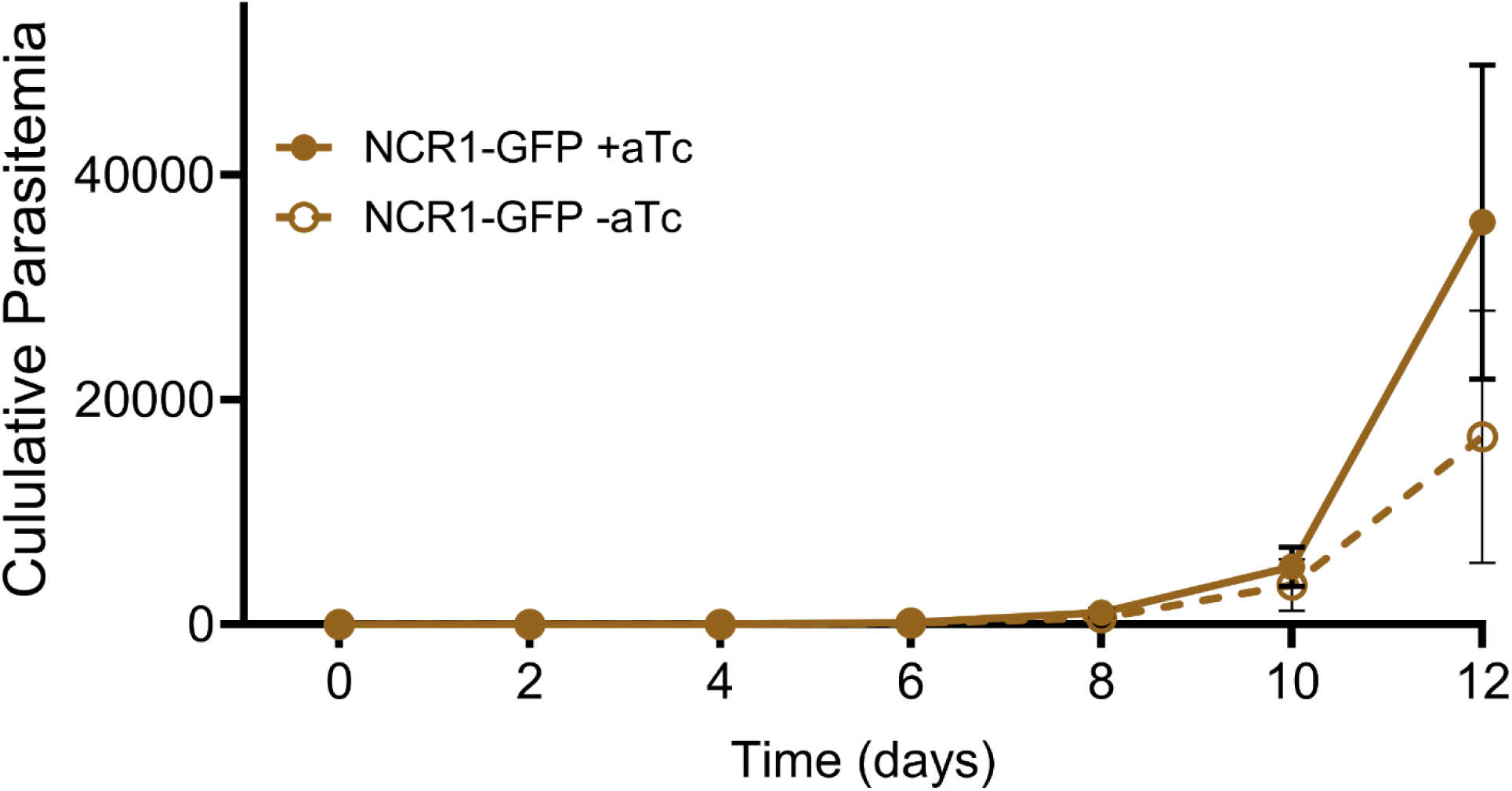
Growth curve of NCR1-GFP +/-aTc. NCR1-GFP parasites were cultured with (+; solid line) or without (-; dashed line) aTc at 1% parasitemia with 2% hematocrit, sub-cultured every 48 hours. Parasite growth was monitored using a flow cytometer. The plot represents cumulative parasitemia calculated by multiplying with the dilution factor over twelve days for two biological replicates.Doubling time +aTc is 0.80 days and -aTc 0.88 days (exponential growth fit on log transformed data in GraphPad prism).

**Fig. S6.**
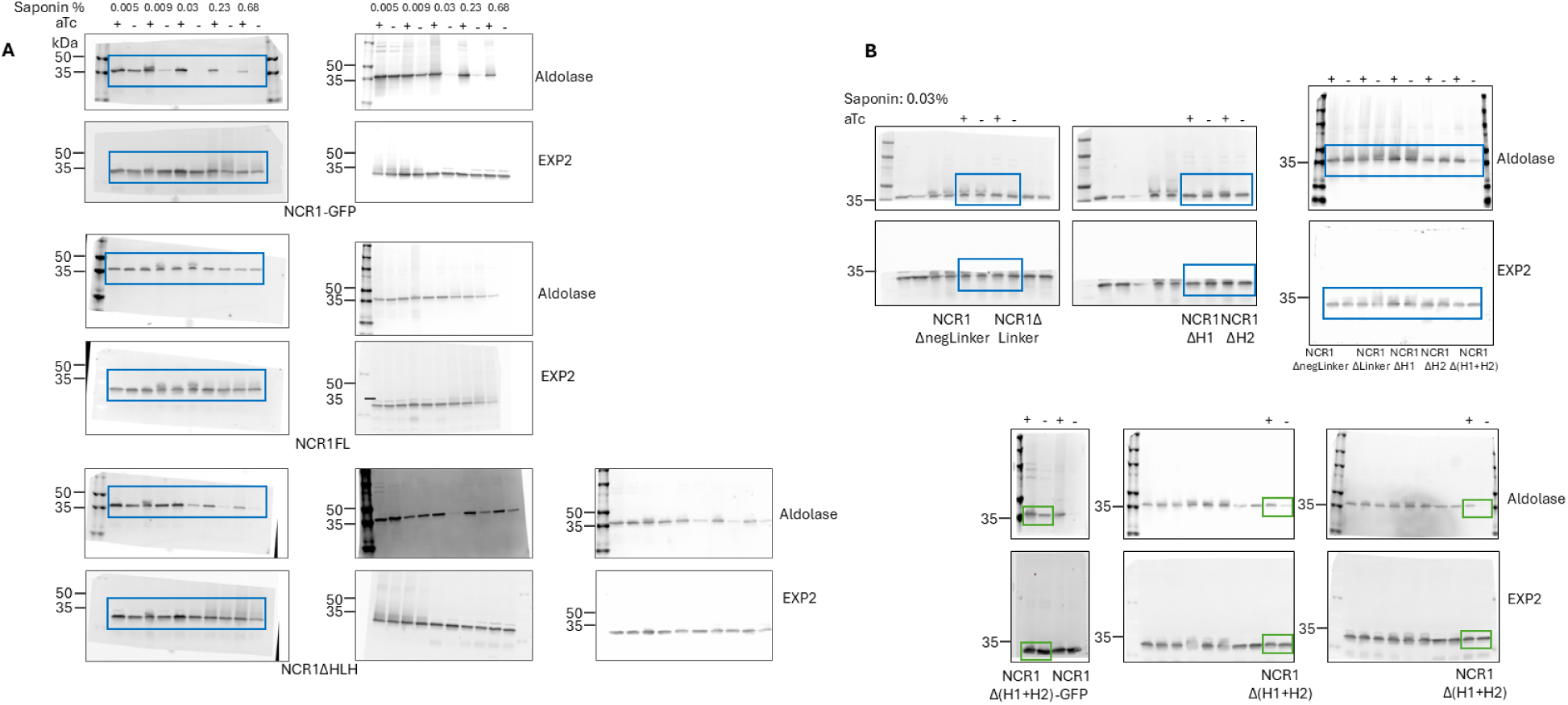
Uncropped western blot images of Figs. 3A and 3C. The blue boxes represent images included in Fig. 3 B and D. The green boxes represent biological and technical replicates for NCR1Δ(H1+H2). The other western blots are biological replicates of individual experiments.

**Fig. S7.**
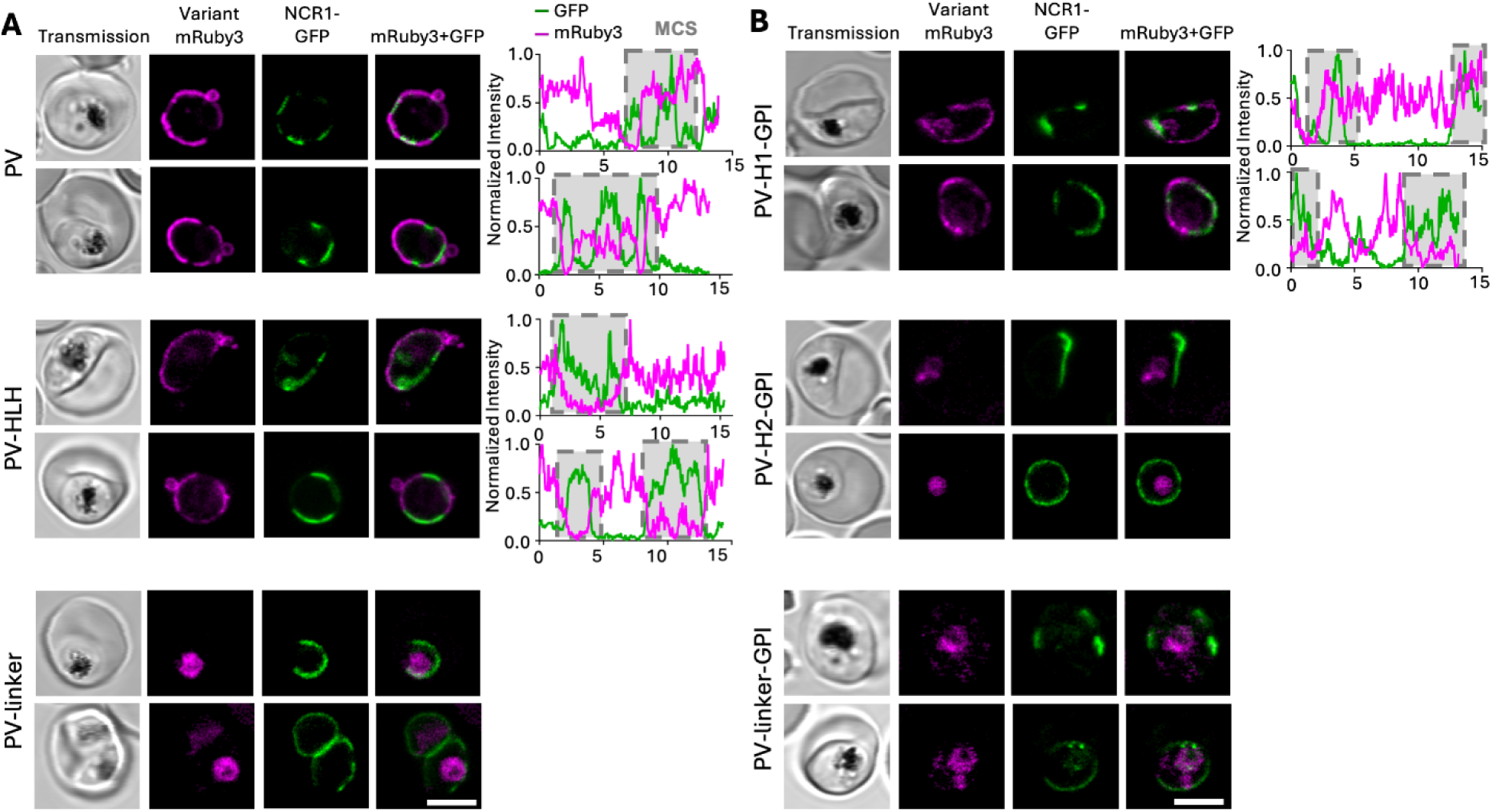
Additional examples of cells from figure 4. A. Confocal microscopy images of parasites expressing NCR1-GFP and PV targeted mRuby3 (PV-mRuby3), HLH-domain-mRuby3 (PV-HLH-mRuby3), and the disordered linker-mRuby3 (PV-Linker-mRuby3). B. Addition of a GPI-anchor to the individual helices H1 and H2: PV-H1-mRuby3-GPI and PV-H2-mRuby3-GPI, and to the linker: PV-linker-mRuby3-GPI. A-B: PV - relatively high: -0.26, low: -0.78; PV-HLH - average: -0.44, low: -0.79; PV-H1-GPI - relatively high: 0.21, low: -0.63. Protein domain diagrams on top of sub-panels introduce PV signal sequence tagged proteins with or without GPI expressed as second copy in the NCR1-GFP parent line. Scale bar: 5 µm.

